# FGF receptor modulates planar cell polarity in the neuroectoderm via Vangl2 tyrosine phosphorylation

**DOI:** 10.1101/2025.05.28.656647

**Authors:** Ilya Chuykin, Sergei Y. Sokol

## Abstract

FGF receptors (FGFR) play pivotal roles in morphogenetic processes including vertebrate neurulation. Planar cell polarity (PCP) signaling coordinates cell polarization in tissue plane and also plays an essential role in neural tube closure. Here we demonstrate abnormal PCP in *Xenopus* neuroectoderm depleted of FGFR1, suggesting a mechanistic connection between FGFR signaling and morphogenesis. FGFR1 associated with the core PCP protein Vangl2 leading to its phosphorylation at N-terminal tyrosine residues. This phosphorylation required FGFR1 activity in frog embryos and mouse embryonic stem cells, extending our observations to mammals. Mutagenesis indicated that the phosphorylation inhibits the interaction of Vangl2 with Prickle and the receptor tyrosine kinase PTK7, leading to the disruption of neuroectodermal PCP. This study identifies a cross-talk between the FGFR and PCP pathways mediated by Vangl2 tyrosine phosphorylation.

## Introduction

The FGF receptor pathway is known to orchestrate cell specification and tissue patterning during embryogenesis. In addition to the regulation of cell fate, cell proliferation and differentiation, FGFR signaling has been implicated in the morphogenetic processes that involve all germ layers ^1–6^. In vertebrates, the pathway consists of eighteen FGF ligands and four receptors from the receptor tyrosine kinase (RTK) superfamily. Upon stimulation, the FGFRs dimerize and the tyrosine kinase domains cross-phosphorylate each other leading to the activation of downstream effectors such as ERK, PI3K, PLCγ, and STAT1 ^7–9^. Although the ERK pathway is considered a principal transducer of the FGFR activity in the embryo, FGFR-dependent developmental signaling has been proposed to depend on additional yet unknown molecular targets. The knocked-in *fgfr1* gene with mutated binding sites for the known mediators of FGF signaling fails to phenocopy the null mutant phenotype in mice, indicating that FGFR1 must possess still uncharacterized modes of signaling ^10,11^.

The planar cell polarity (PCP) pathway coordinates cell alignment in the tissue plane ^12–14^ and is required for many morphogenetic events in vertebrate embryos. Initially discovered in *Drosophila*, ‘core PCP’ components include the transmembrane proteins Van Gogh (Vang), Frizzled (Fz), Flamingo/CELSR, and the cytoplasmic proteins Prickle (Pk) and Dishevelled (Dvl). Due to feedback regulation, Vangl/Pk and Fzd/Dvl complexes segregate to opposite cell boundaries ^15–21^. In vertebrates, Vang-like genes (Vangl) play important roles in neural tube morphogenesis ^22–28^. Mouse studies have demonstrated genetic interactions of *Vangl2* with other genes implicated in PCP, such as *Wnt5*, *Ror2*, *Scribble*, *Celsr1* and the inactive receptor tyrosine kinase *Ptk7* ^29–31^.

In *Xenopus* early embryos, the enrichment of Vangl2 at the anterior surface of each cell is first evident in the posterior neural plate and subsequently extends anteriorly during neurulation, suggesting a posterior origin of the PCP-instructing signal ^32,33^. Because of the posterior expression and activity of several FGF ligands during neurulation ^34–37^, we asked whether the FGFR pathway plays a role in the establishment of PCP. In this study, we uncover a vertebrate-specific tyrosine phosphorylation of Vangl2 in response to FGFR1 and demonstrate a requirement of FGFR signaling for PCP in the *Xenopus* neural plate. Mechanistically, we find that the phosphorylation inhibits the association of Vangl2 with Pk3 and Ptk7, an RTK family pseudokinase that genetically interacts with Vangl2 in neural tube closure ^31,38,39^.

## Results

### Requirement of FGFR1 for neural plate planar polarity

FGF signaling is required for anteroposterior patterning in the neural plate, neural induction ^34,35,40–43^ and neural tube closure ^2^. Whether these functions involve distinct pathway targets and whether a separate signaling branch is responsible for the control of collective cell behaviors remains to be determined.

To investigate a potential link between FGFR1, a commonly expressed FGF receptor ^44,45^, and the core PCP pathway, we examined the distribution of the Vangl2 protein in the plane of the *Xenopus* neural plate of embryos injected with a previously characterized FGFR1 morpholino oligonucleotide (MO) ^46^. We found that injection of FGFR1 MO caused severe neural tube defects that were partially rescued by MO-resistant mouse FGFR1 construct (**Supplementary Fig. 1**). To avoid global effects on neural specification, we generated GFP-marked mosaic clones of FGFR1-morphant cells. Vangl2 was enriched at anterior edges of the uninjected and control MO-injected neuroepithelial cells as reported previously ^32^ (**Fig. 1A,B-B’**). By contrast, the cells containing FGFR1 MO lacked the typical Vangl2 polarity that is characteristic of PCP (**Fig. 1C, D; Supplementary Fig. 2**). We did not observe significant effects of FGFR1 MO on the expression of the pan-neural marker Sox3 suggesting that cell specification was largely intact (**Supplementary Fig. 3**). Inhibition of the anterior enrichment of endogenous Vangl2 (**Supplementary Fig. 4)** and exogenous complexes of Vangl2 and Pk3 ^32,47,48^ (**Supplementary Fig. 5**) has been confirmed using a dominant-negative FGFR1 construct (XFD) ^44^. These observations suggest a role for FGFR1 signaling in establishing PCP in the neural plate.

**Figure 1.**
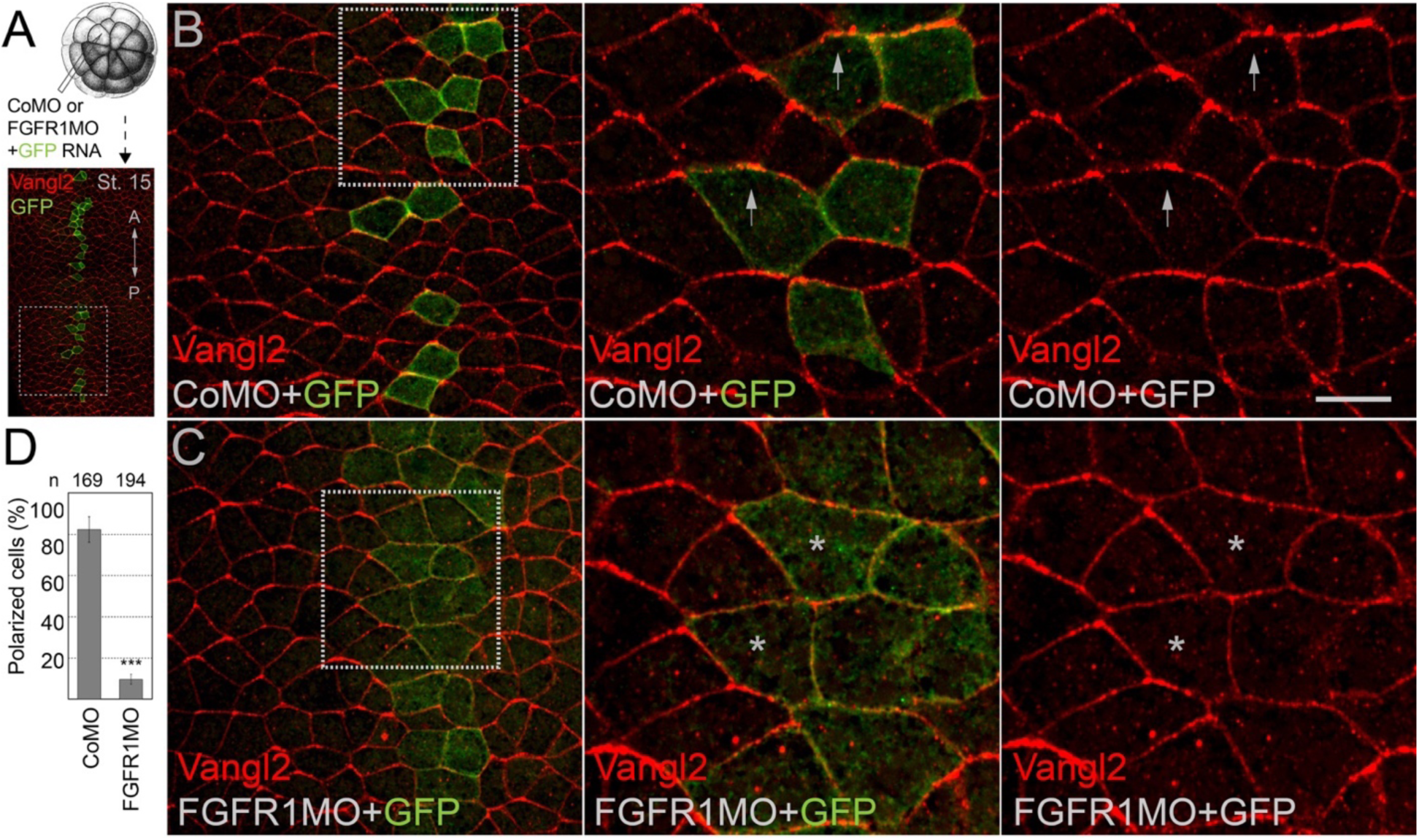
Disrupted planar cell polarity in the neuroectoderm of FGFR1-depleted embryos. (A) Experimental scheme. Thirty-two-cell *Xenopus* embryos were co-injected with 5 nl of control (Co) or FGFR1 morpholino (MO), 15 ng each, along with GFP RNA (90 pg) as a lineage tracer into one dorsal animal blastomere. Embryos were collected at stage 15 (st.15) and co-immunostained for Vangl2 (red) and GFP (green). The anteroposterior (AP) axis is indicated. Representative *en face* neural plate images are shown in (B) and (C), white box areas are magnified on the right to show merged (green+red) and single (red) channel images. (B) Control neuroectoderm with anteriorly polarized Vangl2 (arrows). (C) FGFR1MO-injected neuroectoderm lacking anterior Vangl2 accumulation (asterisks). (D) Frequencies of GFP-positive cells with anteriorly enriched Vangl2 in the CoMO and FGFR1MO injected neuroectoderm. Numbers of scored cells (n) are indicated on the top of each bar. Scoring was performed on 50-90 cells per embryo, three embryos per group. Data represent four independent experiments; *** p<0.001, two-tailed unpaired Student’s *t* test. Scale bar, 30 µm.

In addition to early neuroectoderm, we examined the distribution of Prickle2 (Pk2) that exhibits planar polarization at posterior edges of tailbud skin cells ^49^. We find that FGFR signaling is required for Pk2 planar polarity in the skin (**Supplementary Fig. 6**), suggesting that this pathway may have a broad role in PCP establishment.

### Tyrosine phosphorylation of Vangl2 in response to FGF receptor activation

Although FGF ligand has been previously reported to modulate the polarization of Vangl2 in the developing mouse limb ^50^, the mechanistic link between FGFR signaling and PCP remains unclear. We therefore asked whether endogenous FGFR1 might regulate Vangl2 tyrosine phosphorylation *in vivo*. Notably, the phosphorylation can be detected in untreated ectodermal explants of stage 12 embryos (**Fig. 2A, B**). The signal was reduced by the XFD construct (**Fig. 2B, Supplementary Fig. 7**), by FGFR1 MO (**Fig. 2C**) and after the treatment of the explants with the chemical inhibitor of FGFR activity SU5402 ^51^ **(Supplementary Fig. 8)**, indicating that this phosphorylation event requires endogenous FGFR1 function in *Xenopus* embryos. This conclusion was further supported by experiments using FGFR1/2 double knockout mouse embryonic stem (ES) cells ^52^ (**Fig. 2D,E, F**). The latter result establishes FGFR1/2 as significant contributors to Vangl2 phosphorylation in mouse ES cells, extending our findings to the mammalian model.

**Figure 2.**
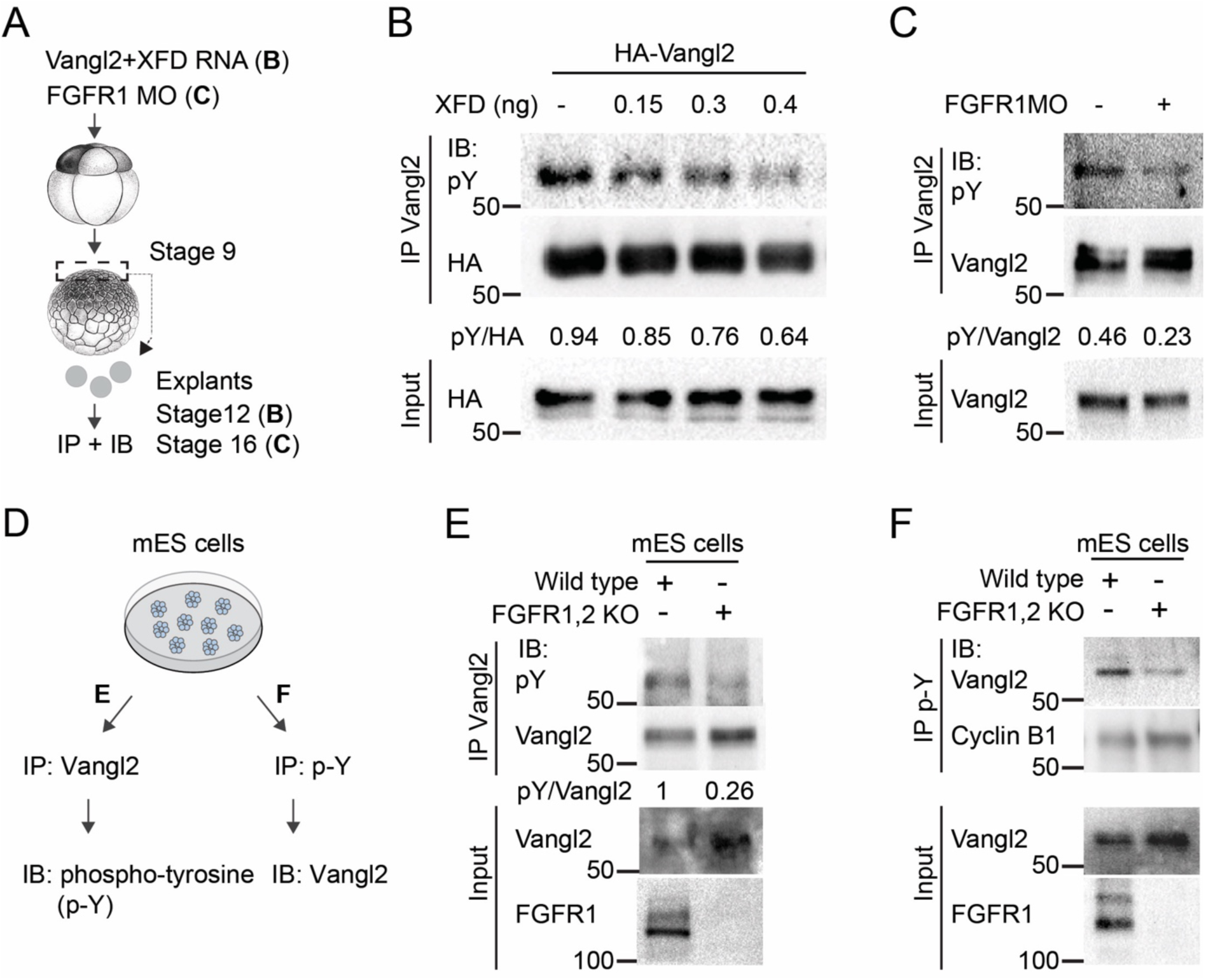
Vangl2 tyrosine phosphorylation requires FGFR signaling. (A-C) FGFR signaling inhibition decreases Vangl2 tyrosine phosphorylation in *Xenopus* embryos. (A) experimental scheme for (B-C). Ectoderm explants were dissected from stage 9 embryos and used for pull down with Vangl2 antibody. (B) Explants were dissected from embryos injected with HA-Vangl2 RNA (40 pg), with or without dominant-interfering FGFR1 (XFD) and cultured until stage 12. (C) Reduction of endogenous Vangl2 tyrosine phosphorylation in ectoderm explants (stage 16) depleted of FGFR1 with FGFR1 MO (20 ng). (D-F) Vangl2 tyrosine phosphorylation depends on FGFR signaling activity in mouse embryonic stem (ES) cells. (D) Experimental scheme. (E, F) Lysates from wild-type and FGFR1/2 double knockout mouse ES cells were precipitated with anti-Vangl2 (E) or anti-phosphotyrosine (pY) (F) antibodies and immunoblotted as indicated. Cyclin B1 is a negative control (F).

To evaluate whether FGFR1 is directly connected to core PCP proteins, we tested whether FGFR1 physically associates with Vangl2. For both exogenous and endogenous proteins, we found that FGFR1 co-precipitates with Vangl2 (**Fig. 3A, Supplementary Fig. 9**), consistent with crosstalk between the two pathways. We next assessed a potential mechanism of PCP modulation by FGFR1. Vangl2 is known to be phosphorylated by Casein kinase I on serine/threonine residues in response to Wnt and Frizzled signaling, leading to reduced gel mobility and altered functional activity ^29,32,53–56^. More recently, tyrosine phosphorylation of Vangl2 has also been reported ^57,58^. We asked whether Vangl2 is phosphorylated at tyrosine residues in embryos expressing exogenous FGFRs. A strong signal was detected by anti-phosphotyrosine (pY) antibody in Vangl2 pulldowns from lysates of embryos co-expressing Vangl2 and FGFR1, FGFR2, FGFR4 and constitutively active FGFR1Y372>C but not those co-expressing Vangl2 and Frizzled3 (**Fig. 3B, Supplementary Fig. 10**). These findings reveal a physical association between FGFR1 and Vangl2 and show that overexpressed FGFRs can stimulate Vangl2 phosphorylation. Together, our findings point to Vangl2 as a molecular target of the FGFR pathway and demonstrate that FGFR is both necessary and sufficient for Vangl2 tyrosine phosphorylation in early embryos.

**Figure 3.**
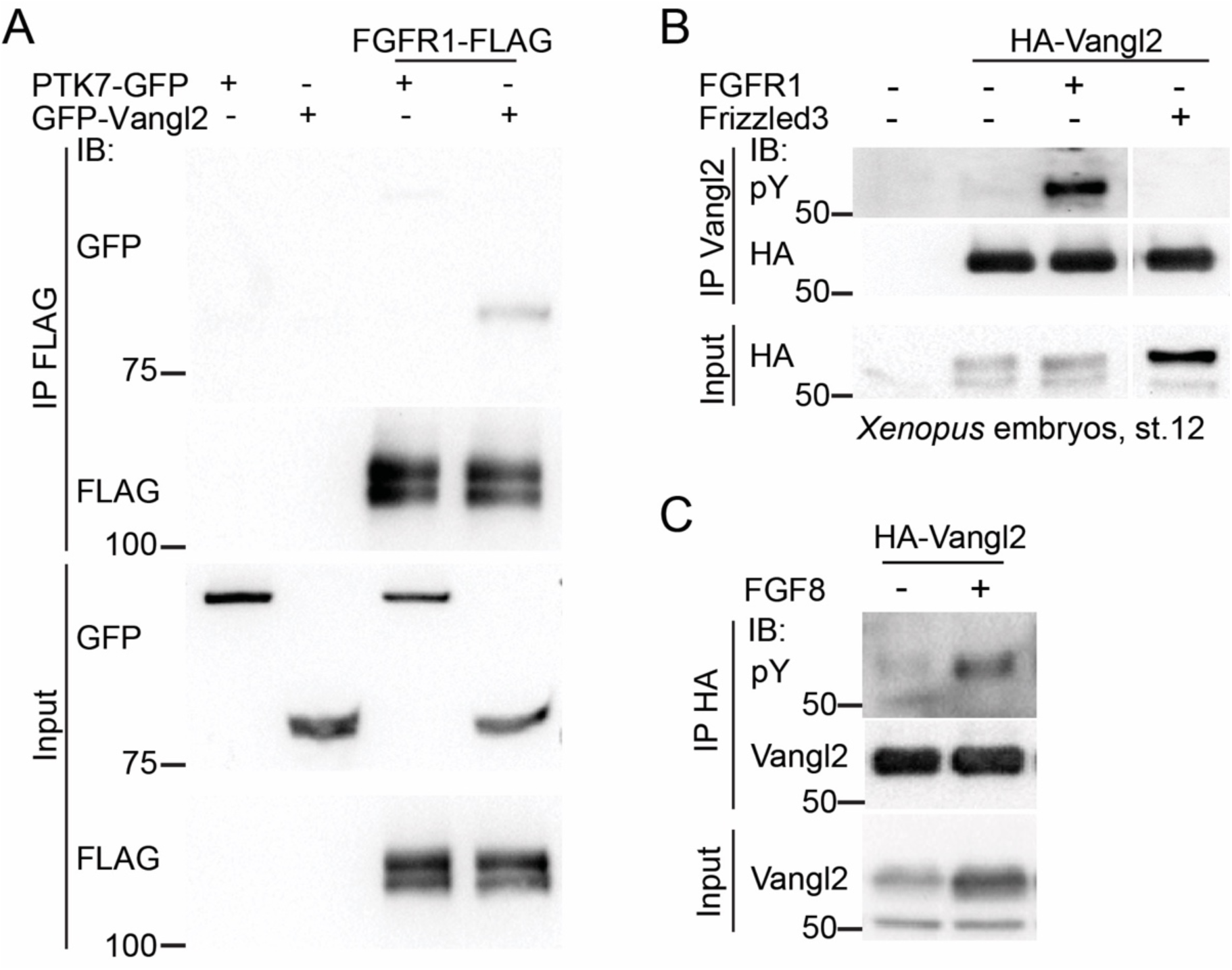
FGFR1 physically associates with Vangl2 and triggers its tyrosine phosphorylation. Four- to eight-cell *Xenopus* embryos were injected with RNAs for GFP-Vangl2, PTK7-GFP and FGFR1-FLAG (100 pg each) (A) or RNAs for HA-Vangl2 (50 pg) and either FGFR1-FLAG (40 pg) or Frizzled3-FLAG (400 pg) (B), lysed at stage 12 and subjected to immunoprecipitation (IP) with anti-FLAG to assess FGFR1-Vangl2 binding (A) or anti-Vangl2 to assess Vangl2 tyrosine phosphorylation (B). PTK7-GFP is a negative control in (A). Frizzled3-dependent Vangl2 band shift in (B) reflects S/T phosphorylation ^54^. An irrelevant portion of the membrane was removed. (C) FGF8 induces Vangl2 phosphorylation in *Xenopus* embryos. Embryos were injected with HA-Vangl2 with or without FGF8 plasmid DNAs (50 pg) each, lysed at stage 14 and precipitated with HA-trap followed by immunoblot with anti-pY antibodies.

### Lack of correlation between ERK activation and Vangl2 phosphorylation and a role for FGF ligands

An important question is whether FGFR signals to Vangl2 via a mechanism distinct from its signaling to the well-known mediator ERK ^7^. Another critical issue is which FGF ligand activates Vangl2 phosphorylation in response to FGFRs. We compared the induction of Vangl2 tyrosine phosphorylation and phospho-ERK1/2 (active ERK), in ectodermal explants in response to FGFR and FGF2. FGFR1 triggered a dose-dependent increase in Vangl2 tyrosine phosphorylation, whereas only a weak synergy was observed between FGF2 and FGFR1, and FGF2 alone had no effect (**Supplementary Fig. 8, Supplementary Fig. 11**). By contrast, FGF2 strongly synergized with FGFR1 in upregulating phospho-ERK1/2. Both phosphorylation events were sensitive to SU5402, indicating dependence on FGFR activity (**Supplementary Fig. 11**).

While searching for a potential inducer of Vangl2 tyrosine phosphorylation among the FGF ligands present during neurulation, we found that FGF8 can stimulate Vangl2 tyrosine phosphorylation (**Fig. 3C**). Since FGF8 is expressed in the posterior neural plate ^36,37^, it is a good candidate for the ligand stimulating FGFR1 signaling to Vangl2.

We next tested FGFR1^FCPG^ a mutated FGFR1 construct lacking the binding sites for known signaling intermediates and deficient in the ability to activate most of the known FGFR1 signaling pathways ^10^. FGFR1^FCPG^ did not activate ERK, consistent with a previous study ^10^, but induced Vangl2 tyrosine phosphorylation even more efficiently than wild-type FGFR1 (**Fig. 4**). This observation strengthens our hypothesis that Vangl2 is phosphorylated via a noncanonical pathway.

**Figure 4.**
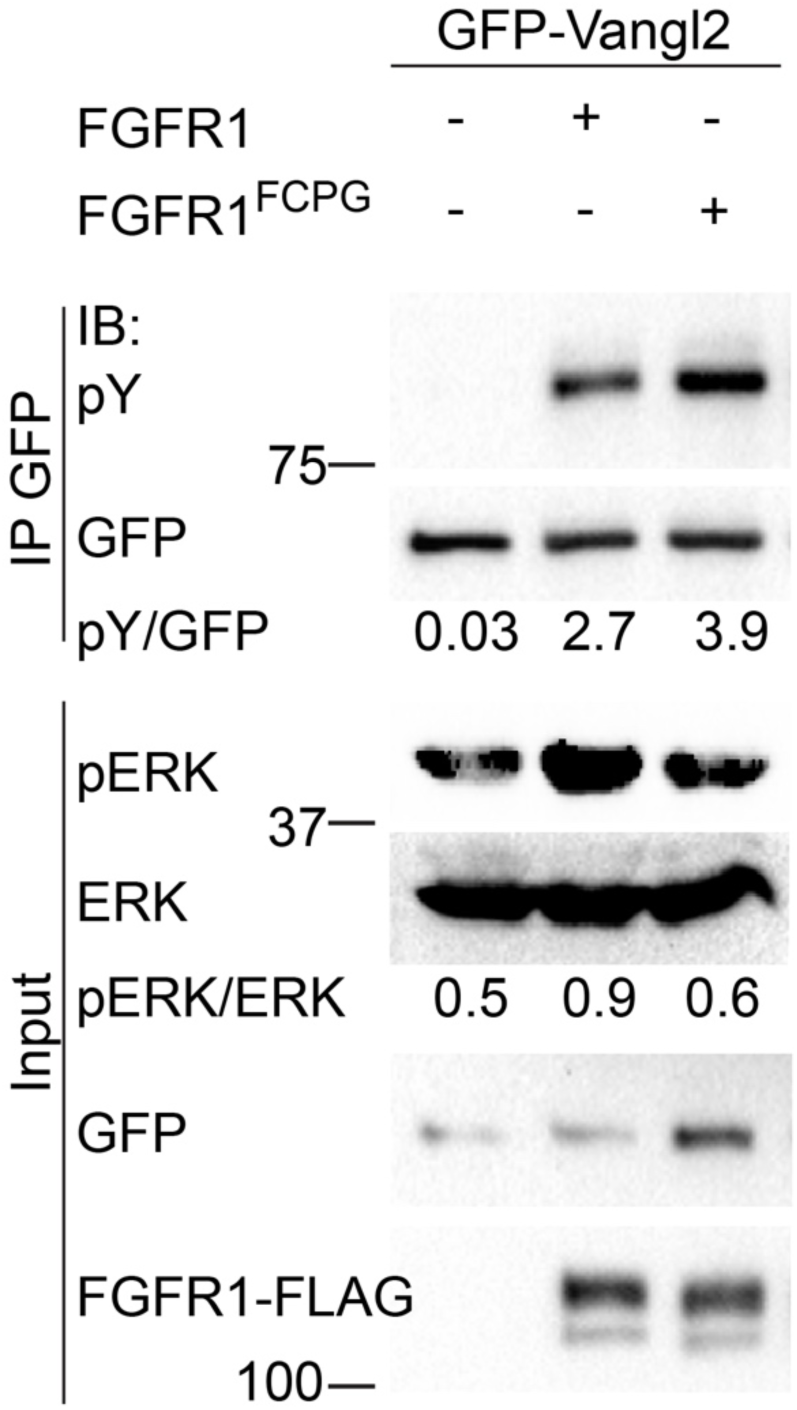
Vangl2 tyrosine phosphorylation by FGFR1 containing mutations that prevent ERK activation. Four-to-eight cell embryos were injected into four animal blastomeres with GFP-Vangl2 RNA, FGFR1-FLAG or FGFR1^FCPG^-FLAG RNA (40 pg each), and were cultured until stage 12. GFP-Vangl2 was immunoprecipitated (IP) from embryo lysates using GFP-trap beads followed by immunoblotting (IB) with anti-pY, anti-GFP, anti-FLAG, anti-pERK1/2 and anti-ERK1/2 antibodies. FGFR1^FCPG^ containing mutations in the binding sites of known signaling mediators ^10^ induced Vangl2 tyrosine phosphorylation but did not upregulate ERK1/2 activity.

### Vangl2 N-terminal tyrosine phosphorylation inhibits planar polarity

To identify specific Vangl2 amino acid residues that become phosphorylated in response to FGFR1, we replaced tyrosine residues with phenylalanines. RNAs encoding FGFR1 and Vangl2 mutants with Y>F substitutions were co-injected into *Xenopus* embryos and the phosphorylation was analyzed in HA-Vangl2 pulldowns using anti-pY antibody. The triple substitution of Y7, Y10, Y12 to phenylalanine strongly decreased FGFR1-induced Vangl2 tyrosine phosphorylation, while the Y341, Y342, Y343 substitutions had only a minor effect (**Fig. 5A)**. Further mapping confirmed that Y10 and Y12 are essential and that the ‘kinase-dead’ form of FGFR1 (D623>A) is completely inactive in this assay (**Supplementary Fig. 12**). Notably, the equivalent of the tyrosine phosphorylation site identified in *Drosophila* Van Gogh ^57^ was not phosphorylated in this experiment (Y308, **Supplementary Fig. 12**). Our findings indicate that the N-terminal tyrosines Y10 and Y12 are the dominant sites in Vangl2 that are phosphorylated in response to FGFR1. The alignment of the N-terminal amino acid sequence shows that the identified sites are conserved among vertebrates (**Fig. 5B**).

**Figure 5.**
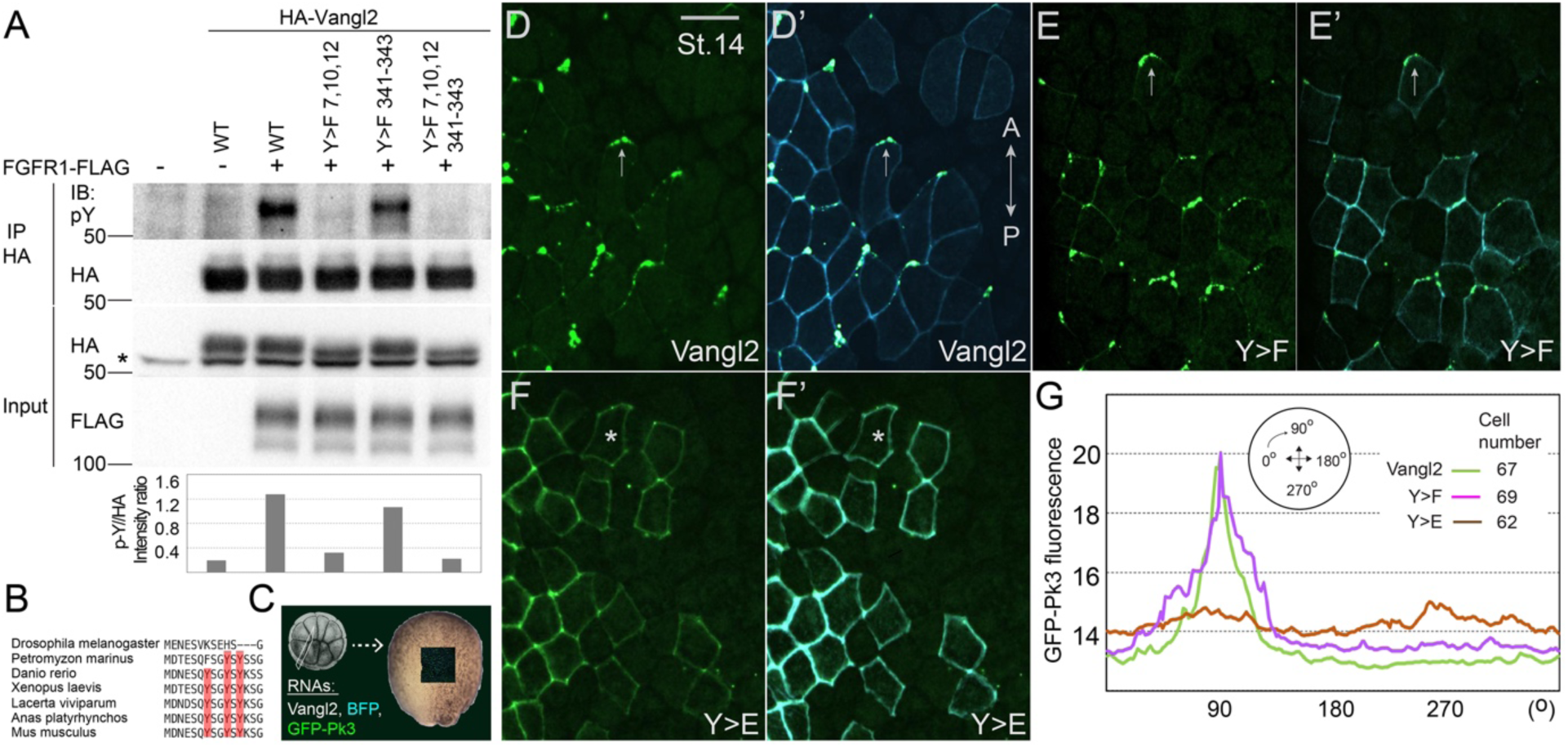
Anterior accumulation of the Vangl2/Pk3 complex in the neuroectoderm is inhibited by N-terminal tyrosine phosphorylation of Vangl2. (A) Mapping major phospho-tyrosine (pY) sites in Vangl2. Embryos were injected with 40 pg of RNA encoding FGFR1 and the indicated HA-Vangl2 constructs. Vangl2 phosphorylation was analyzed in anti-HA pull-downs from stage 13 embryo lysates. Immunoblotting (IB) was performed using anti-pY, anti-HA or anti-FLAG antibodies, as indicated. One representative set of duplicate samples is shown. The graph below shows average pY/HA intensity ratios for the duplicates. (B) Alignment of N-terminal amino acid sequences of Vangl2 from several chordate species and *Drosophila* Van Gogh. The N-terminal tyrosine cluster (in red) is conserved in vertebrates but not in *Drosophila*. (C-E) Planar polarization of Vangl2 tyrosine phosphosite mutants in the neuroectoderm. (C) Experimental scheme. Two dorsal blastomeres of 16-cell embryos were coinjected with myrBFP RNA (80 pg) and RNAs encoding GFP-Pk3 (150 pg) HA-tagged Vangl2 (in D), Y7,10,12>F (Y>F, in E) or Y7,10,12>E (Y>E, in F) (20 pg each). Embryos were fixed at stages 14-15, and GFP and BFP fluorescence were imaged. Anterior enrichment of GFP-Pk3 is indicated by arrows (D-E’), while a non-polarized cell is marked by an asterisk (F-F’). The anteroposterior (AP) axis is indicated. (G) Quantification of GFP fluorescence for the GFP-Pk3-Vangl2 complexes in mosaically expressing cells is shown as a graph. Mean fluorescence is plotted along the cell circumference as a function of circular angle from 0 to 360 degrees relative to the AP axis, with fluorescence intensity shown in green (HA-Vangl2), pink (HA-Vangl2Y>F), and brown (HA-Vangl2Y>E) lines. The number of scored cells is indicated. The data are representative of four independent experiments.

We next asked whether the observed Vangl2 tyrosine phosphorylation may influence the phosphorylation at previously reported Cluster 1 and Cluster 2 serine/threonine sites ^29^. Using antibodies specific for phospho-T78,S79,S82 and phospho-S14,S17 peptides, we observed reduced phospho-S14,S17-Vangl2 levels (Cluster 2) in the Y7,10,12>F Vangl2 construct whereas the phosphorylation at the Cluster 1 was not affected (**Supplementary Fig. 13**). This finding indicates that tyrosine phosphorylation of Vangl2 promotes serine phosphorylation in the adjacent Cluster 2 sites.

Our subsequent experiments focused on the physiological significance of posttranslational modifications at these sites. To assess the ability of unphosphorylatable or phosphomimetic Vangl2 proteins to associate with other PCP proteins, we studied their interactions with Prickle, a known component of the anterior PCP complex ^32,47,48^. RNAs encoding Pk3 and mutated Vangl2 constructs were co-injected into dorsal blastomeres at the 16-cell stage at doses that did not affect normal morphogenesis (**Fig. 5C**). Both wild-type and nonphosphorylatable Vangl2 constructs formed anterior complexes with Pk3, whereas no anterior accumulation was observed with phosphomimetic Vangl2 (**Fig. 5D-G**).

We also tested whether Vangl tyrosine phosphorylation regulates the Vangl2-Pk3 association. This was carried out using a modified proximity biotinylation assay ^59^. We observed reduced interaction of phosphomimetic Vangl2 with Pk3 as compared to the wild-type and nonphosphorylatable Vangl2 constructs (**Fig. 6).** These results suggest that Vangl2 tyrosine phosphorylation decreases the amount of the Vangl2-Pk3 complex, likely reducing the signaling activity of Vangl2.

**Figure 6.**
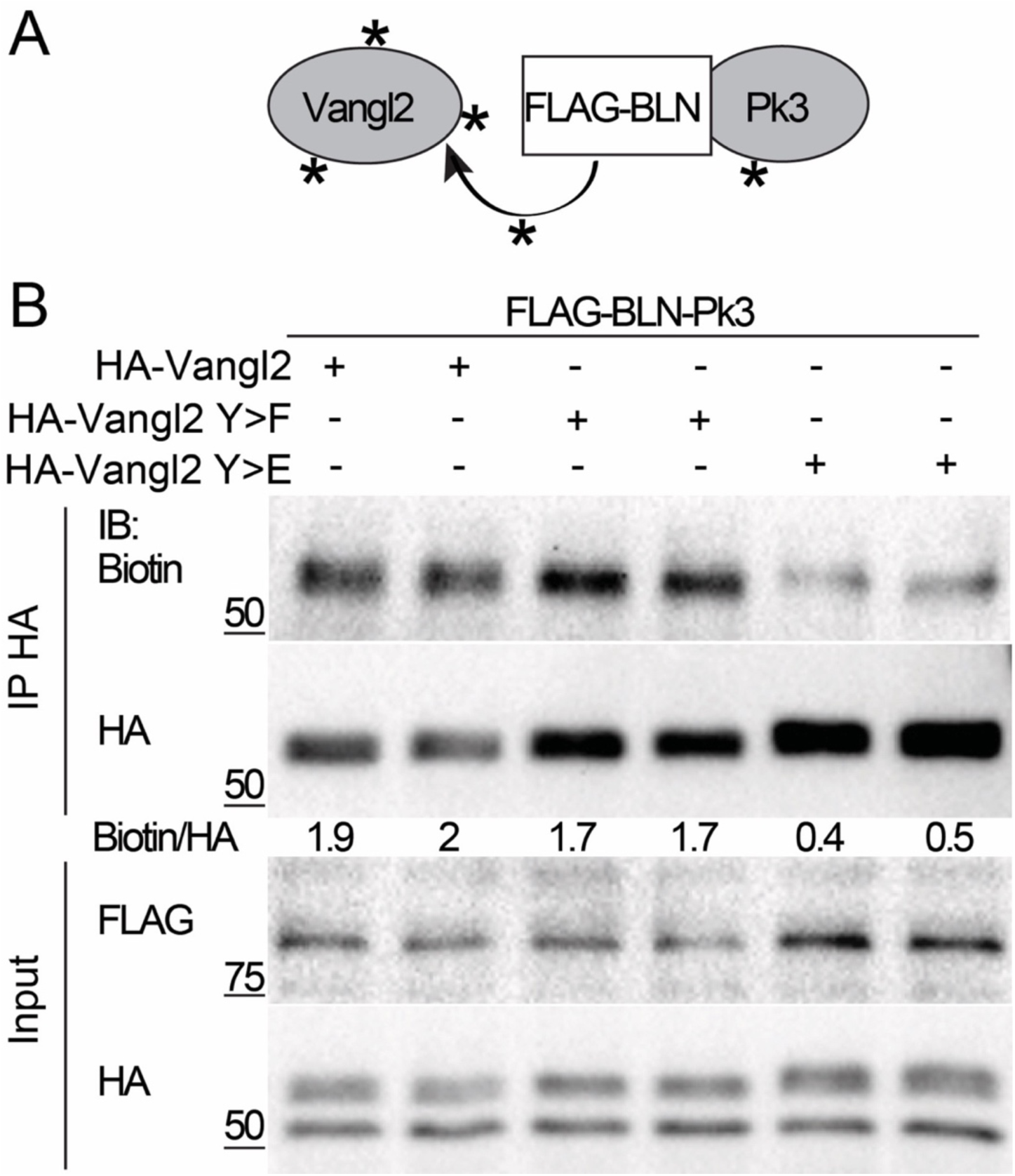
The Vangl2-Prickle3 interaction is inhibited by Vangl2 tyrosine phosphorylation. (A) Schematic for the proximity biotinylation assay to assess the interaction between Vangl2 and Prickle3 (Pk3). Vangl2 is biotinylated (asterisks) when in proximity to the Pk3 fused to the large N-terminal fragment of a bacterial biotin ligase (BLN). (B) Animal blastomeres of four-to-eight-cell stage embryos were co-injected with 100 pg of FLAG-BLN-Pk3 RNA and 40 pg of HA-Vangl2, HA-Vangl2 Y7,10,12>F (Y>F), or HA-Vangl2 Y7,10,12>E (Y>E) RNAs. At stages 8-9, 20 nl of biotin (0.8 mM) was injected into the blastocoel. Embryos were collected at stage 13, and Vangl2 constructs were pulled down using anti-HA antibody. Biotinylation of Vangl2 and protein expression levels were assessed in independent duplicate samples using anti-biotin, anti-HA, and anti-FLAG antibodies. Data represent three independent experiments.

The significance of Vangl2 tyrosine phosphorylation for PCP has been expanded to the complex of Vangl2 and the receptor tyrosine kinase PTK7, a transmembrane protein that genetically interacts with Vangl2 ^31^ and functions in PCP via an unknown mechanism ^31,38,39,60^. We observed that wild-type Vangl2 co-precipitated with PTK7 in lysates of stage 12 *Xenopus* embryos (**Fig. 7A, B**). Notably, this complex formed more efficiently with nonphosphorylatable Vangl2, whereas it was strongly reduced with the phosphomimetic Vangl2 construct (**Fig. 7A, B**), arguing that Vangl2 phosphorylation may control the Vangl2-PTK7 association.

**Figure 7.**
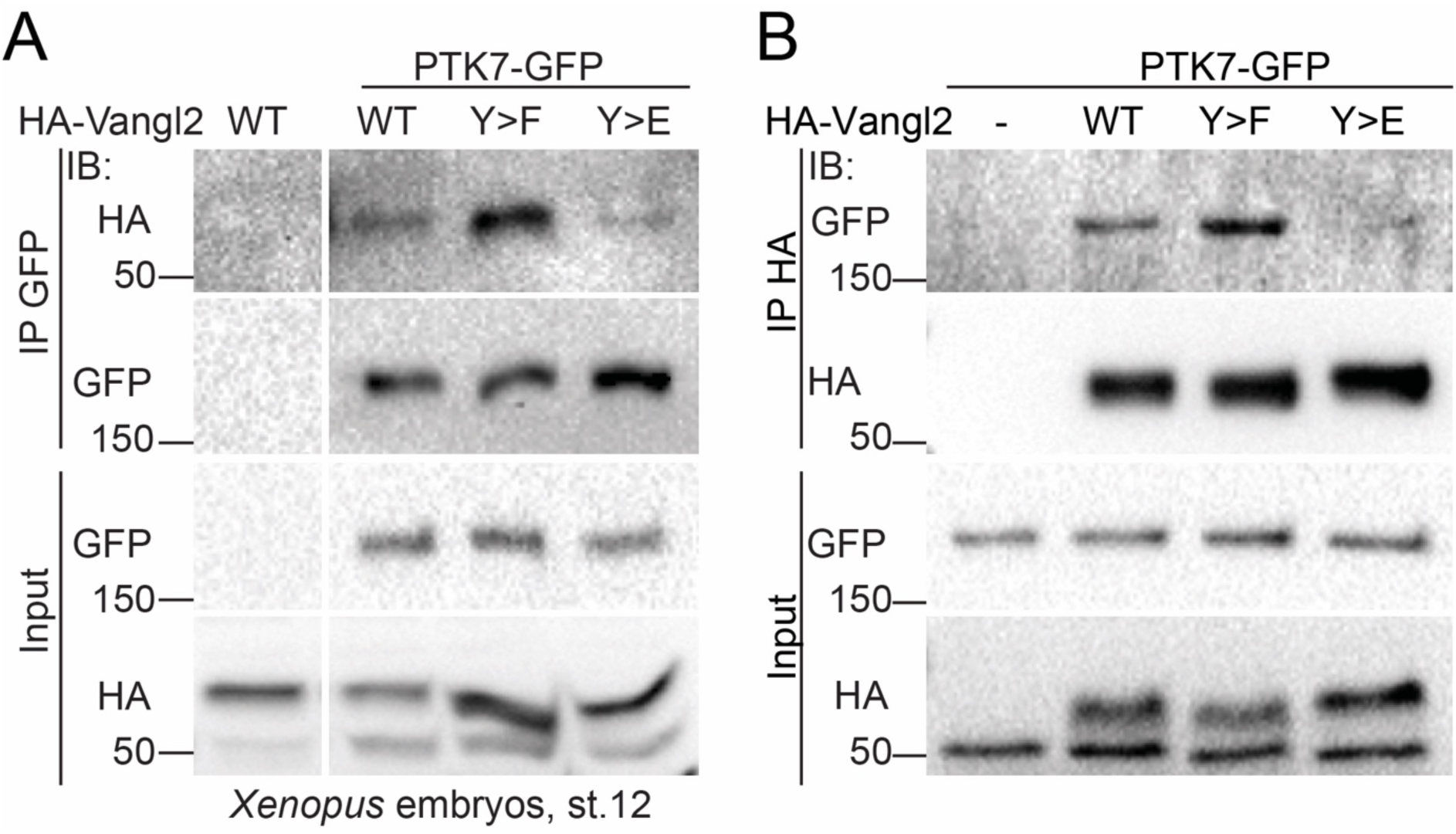
Vangl2 forms a complex with PTK7 in a tyrosine phosphorylation-dependent manner. (A, B) Vangl2-PTK7 binding is enhanced by non-phosphorylated Vangl2 and reduced by phosphorylated Vangl2. Four-to-eight-cell *Xenopus* embryos were injected with RNAs encoding HA-Vangl2, HA-Vangl2 Y7Y10Y12>F (Y>F) or HA-Vangl2 Y7Y10Y12>E (Y>E) (40 pg each) as indicated, with or without PTK7-GFP RNA, 100 pg, into four animal blastomeres. Embryos were collected at stage 12 and PTK7-GFP (A) or HA-Vangl2 (B) were immunoprecipitated using GFP-trap or anti-HA antibody, respectively, for Vangl2 analysis (A) or PTK7 analysis (B). Protein levels in pull-downs and lysates were analyzed with anti-GFP and anti-HA antibodies. Irrelevant part of the membrane was removed (A). Data are representative of three experiments.

To address the significance of this observation, we asked whether PTK7 can functionally synergize with Vangl2 in the induction of neural tube defects. Notably, strong synergy was observed when PTK7 was coexpressed with nonphosphorylatable Vangl2 but not phosphomimetic Vangl2, pointing to the physiological role of Vangl2-PTK7 complex in the neural plate morphogenesis (**Fig. 8A-D**). Together, these experiments highlight a role of the Vangl2-PTK7 complex in PCP.

**Figure 8.**
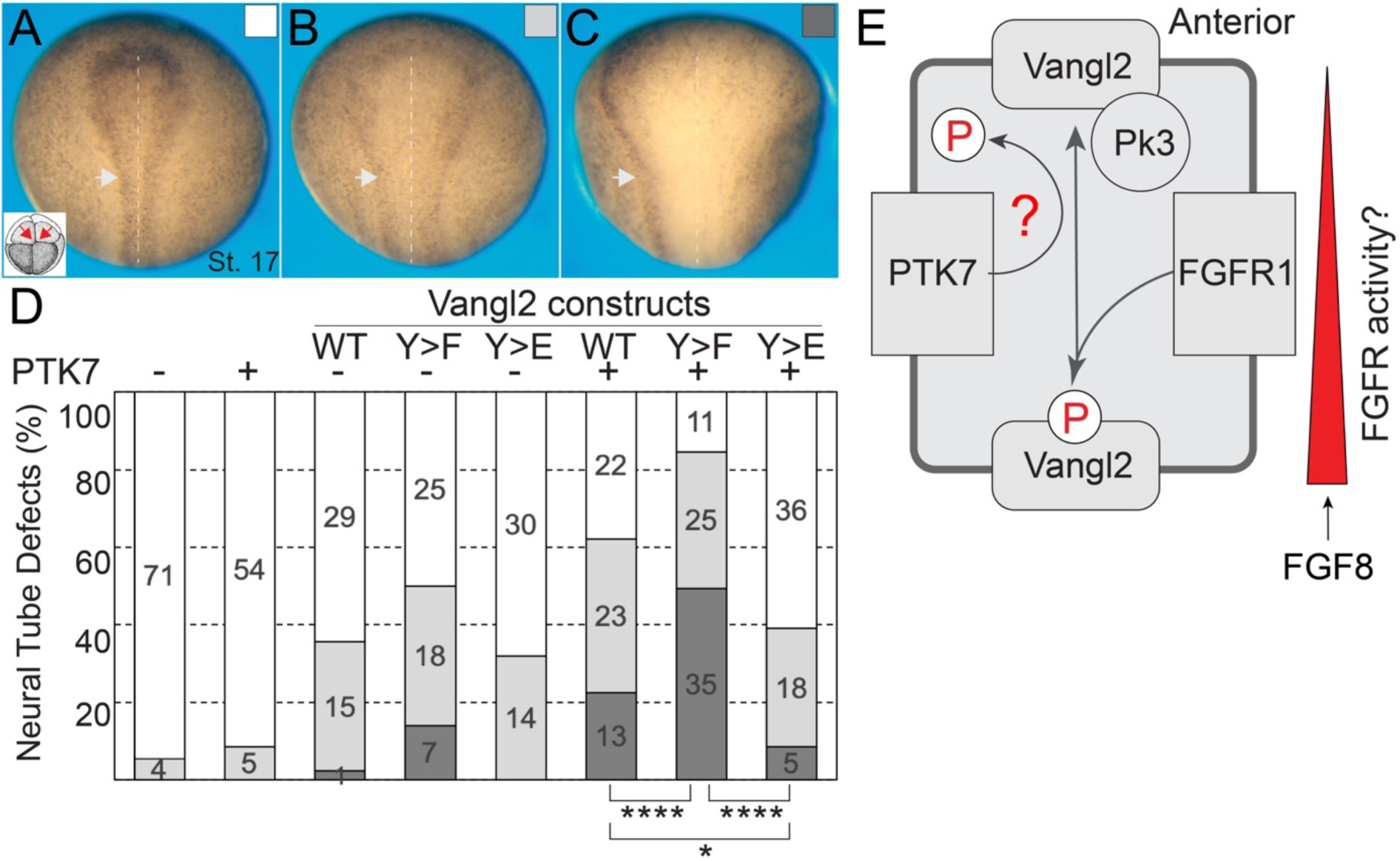
Synergistic effects of Vangl2 and PTK7 on neural tube closure depend on Vangl2 tyrosine phosphorylation. Neural folding defects were assessed in embryos injected with RNAs encoding PTK7 and different forms of Vangl2 (200 pg each) into two dorsal animal blastomeres (inset in A, red arrows). Dorsal views of representative embryos exhibit no (A), mild (B) or severe (C) disruption of neural tube closure. Co-injection of PTK7 and Y>F Vangl2 RNAs causes stronger neural tube defects (NTDs) compared to the co-injection of the wild type or Y>E Vangl2 RNAs. The degree of neural tube closure was scored at stage 17 by the distance between the opposing neural folds near the brain-spinal cord border (arrowheads), anterior is to the top. Dashed line marks the midline. (D) Quantification showing frequencies of normal neural folds (A), as compared to mild (B) and severe (C) defects. The number of scored embryos is shown in the graph. The significance was determined by full 2×3 Fisher’s exact test that was calculated in Prism 10 software; **** p<0.0001, * p<0.05. Data represent two independent experiments. (E) Model. Graded FGF/FGFR1 activity triggers Vangl2 tyrosine phosphorylation at posterior cell edges, inhibiting the formation of Vangl2-Pk3 and Vangl2-PTK7 complexes, leading to reduced posterior accumulation of Vangl2. PTK7 is proposed to act as a feedback regulator and FGFR1 antagonist, promoting Vangl2 dephosphorylation through phosphatase recruitment.

## Discussion

This work demonstrates that FGFR1 physically associates with Vangl2 and triggers its tyrosine phosphorylation. The lack of correlation between Vangl2 phosphorylation and ERK activation points to distinct signaling branches downstream of FGFR1. We propose that FGFR1 tyrosine kinase that is activated by an FGF ligand, such as posteriorly expressed FGF8, binds and directly phosphorylates Vangl2 in a spatially restricted manner (**Fig. 8E**). In the absence of additional evidence, we cannot formally exclude the involvement of other protein tyrosine kinases. It is also possible that the same Vangl2 tyrosine phosphorylation sites serve as substrates for different RTKs expressed in various embryonic tissues ^58^. The requirement of FGFR1/2 signaling for Vangl2 phosphorylation in *Xenopus* embryos and mouse ES cells and the demonstration of its importance for PCP in both the neuroectoderm and the tadpole skin underscore the conservation of this signaling event in different embryonic tissues in vertebrates.

The anterior enrichment of endogenous Vangl2 protein is lost in the FGFR1-depleted neuroectoderm, suggesting an important role of FGFR1 in PCP. We found that the interaction of Pk3 with phosphomimetic Vangl2 assessed by proximity biotinylation was reduced and the complex of Pk3 with phosphomimetic Vangl2 was not anteriorly polarized. These findings are consistent with N-terminal tyrosine phosphorylation preventing the formation of Vangl2-containing PCP complexes at the posterior edges of neuroepithelial cells (**Fig. 8E**). Our finding that phospho-S14,S17-Vangl2 levels are reduced in the phosphomutant Vangl2 construct indicates that the phosphorylation of Vangl2 at Cluster 2 residues mediates cross-talk between the FGF and PCP pathways during embryonic patterning and morphogenesis ^29,55,56^. Future studies will define other biological processes affected by FGFR-dependent Vangl2 phosphorylation, including Vangl2 membrane trafficking ^55,61^, lipidation ^62^ or the regulation of Vangl2 degradation ^63,64^.

Surprisingly, the anterior enrichment of Pk3 in the neuroepithelial cells was preserved in complex with the nonphosphorylatable form of Vangl2, possibly due to limitations of our assay. Although the nonphosphorylatable Vangl2 behaved similarly to the wild-type when tested for the planar polarization of anterior Vangl2-Pk3 complexes, we discovered that it strongly bound PTK7, another planar polarity component. By contrast, the association of PTK7 with phosphomimetic Vangl2 was barely detectable, indicating that Vangl2 binds PTK7 in the phosphorylation-dependent manner.

Our findings extend previous reports of the genetic interactions between Vangl2 and PTK7 during mammalian neural tube closure ^31^, and point to a mechanistic role of PTK7 in PCP regulation. Currently it remains unknown how PTK7 and Vangl2 modulate each other. Importantly, the synergistic effect of Vangl2 and PTK7 on neural tube closure correlated with the phosphorylation status of N-terminal tyrosines in Vangl2, underscoring the physiological significance of this phosphorylation. Searching protein interaction databases such as BioGRID, we found multiple protein tyrosine phosphatases associated with PTK7. We propose that PTK7 functions in PCP as a feedback regulator, because it can facilitate Vangl2 phosphate removal by PTK7-associated phosphatases or compete with FGFR for Vangl2 binding. Additional loss-of-function experiments are needed to understand the function of PTK7 in PCP.

## METHODS

### Plasmids and mRNA synthesis

Plasmids encoding *Xenopus* GFP- and HA-tagged Vangl2 ^48^, mouse FGFR1-3xFLAG, FGFR1^FCPG^-3xFLAG ^10^ and FGFR2-3xFLAG ^52^, human FGFR1, FGFR1Y372>C and FGFR4 ^65^, dominant-interfering-FGFR1 (XFD) ^44^, GFP ^32^, mCherry ^59^, Fz3-FLAG ^66^, FLAG-BLN-tagged Pk3 ^59^, pCS107-GFP-Pk2 ^49^, pCS2-FGF8 ^37^ have been described previously. pCS2-PTK7-GFP was a gift of Annette Borchers. The pCS2-myristoylated BFP-HA and the corrected GFP-Pk3 construct ^48^, fitting the predicted protein sequence, were gifts of Miho Matsuda. HA-tagged Y>F and Y>E point mutants of Vangl2 and kinase-dead FGFR1D623>A mutant ^67^ were generated using gBlock fragments (Integrated DNA Technologies). pCS2-XFD-3xFLAG construct was generated by PCR. Capped mRNAs were synthesized using mMessage mMachine kit (Ambion, Austin, TX).

### *Xenopus* embryo culture and microinjections

The study was conducted in strict accordance with the recommendations outlined in the Guide for the Care and Use of Laboratory Animals by the National Institutes of Health. The animal protocol received approval from the Institutional Animal Care and Use Committee (IACUC) at the Icahn School of Medicine at Mount Sinai. *Xenopus laevis* eggs were in vitro fertilized and cultured in 0.1x Marc’s Modified Ringer’s solution (MMR) ^54^. For microinjections, 4-16 cell embryos were transferred to a 3% Ficoll 400 solution (GE Healthcare) in 0.6x MMR and were injected with 5-10 nl of a solution containing mRNAs or morpholino. To achieve mosaic expression of PCP complexes or FGFR1-depletion in the neural plate the embryos were injected into two dorsal blastomeres of 16-32-cell embryos. The quantities of injected mRNAs were optimized in preliminary dose-response experiments, and the specific amounts are indicated in the figure legends.

### Immunostaining, fluorescent protein detection, imaging and quantification

To identify PCP complexes in the neural plate, embryos were collected at stage 14, and the vitelline membrane was manually removed. RNAs encoding GFP-Pk3, HA-Vangl2 wild type and mutated Y7Y10Y12>F or Y7Y10Y12>E constructs and mBFP were co-injected at the established doses (as specified in the figure legends), which did not have impact on normal development as was established in preliminary experiments. Embryos were fixed in MEMFA solution for 40 min. Neural plate explants were then dissected, mounted in the Vectashield mounting medium (Vector) and GFP-Pk3 and mBFP fluorescence was detected and scored.

For detection of endogenous Vangl2, embryos were collected at stages 15-16, fixed in 2% TCA for 30 min, and stained with with the rabbit polyclonal anti-Vangl2 antibody (1:100) ^32^. Sox3 was detected with mouse monoclonal Sox3 antibody (1:100; DSHB, DA5H6). For tracing FGFR1-depleted cells, embryos were co-injected with 15 ng of FGFR1 MO and 90 pg of GFP RNA. Mouse monoclonal (Santa Cruz, GFP-B2) or rabbit polyclonal (Invitrogen, A6544) anti-GFP antibodies were used for co-immunostaining at 1:100 and 1:500 dilutions, respectively.

Secondary antibodies were against mouse or rabbit IgG conjugated to Cy2 or Cy3 (1:500, Jackson ImmunoResearch). Standard specificity controls were conducted to verify the absence of cross-reactivity and to ensure that no staining occurred in the absence of primary antibodies.

Images of whole neural plate explants were acquired using tiling and subsequent stitching at the BC43 (Andor) confocal microscope. The quantification of planar polarity of the HA-Vangl2 constructs in complexes with GFP-Pk3 complexes (Vangl2, Vangl2 Y7Y10Y12 >F or >E constructs) was performed using the Fiji software with the Azimuthal Average plugin available at https://imagej.nih.gov/ij/plugins/azimuthal-average.html, following the described steps ^54^. BFP fluorescence was used to identify cell boundaries in mosaic cells, which were selected using the circle tool in Fiji. The fluorescence intensity of GFP-Pk3 was quantified along the radius divided to 180 bins within cells expressing PCP complexes.

To determine the effect of FGFR1MO (15 ng) on neural cell fate, Sox3 immunofluorescence was measured in 3 × 10⁴ µm² rectangular regions of interest on the FGFR1MO-injected (GFP-marked) and uninjected sides of the neural plate in three independent embryos.

### Preparation of ectodermal explants, proximity biotinylation

Four-to-eight-cell embryos were injected with RNAs or FGFR1 MO, as indicated in figure legends, vitelline membrane was removed at stage 9 and ectoderm explants were dissected and cultured with or without of 25-100 ng/ml of FGF2 with or without 100 µM SU5402 (Millipore Sigma) until siblings reached stage 12 or stage 16 when they were collected for immunoprecipitation and immunoblotting analysis.

For proximity biotinylation ^59^ ^68^, embryos were injected into the animal pole of four-to-eight-cell embryos with RNAs encoding FLAG-BLN-Pk3, 100 pg, and HA-Vangl2 constructs, 40 pg. The embryos were allowed to develop until stage 8-9, followed by injection of 20 nl of a 0.8 mM biotin (Millipore Sigma) solution into the blastocoel. Embryos were lysed and the protein biotinylation was assessed in embryo lysates and pulldowns obtained with mouse anti-HA (12CA5) antibody. Stages are indicated in the figure legends. Immunoblotting was carried out with goat anti-biotin-HRP antibodies (Cell Signaling) or goat anti-biotin antibody (Pierce), anti-FLAG (M2, Millipore Sigma), rabbit anti-HA (Bethyl) antibodies, mouse anti-HA (12A5), rabbit anti-phospho-ERK (Cell signaling), rabbit anti-ERK (Cell Signaling), mouse anti-Vangl2 (C8, Santa Cruz), rat monoclonal anti-Vangl2 (2G4, Millipore Sigma), mouse anti-GFP (B2, Santa Cruz). Chemiluminescence was captured by ChemiDoc (BioRad) and the Biotin/HA chemiluminescence intensity ratios were quantified using the ChemiDoc software.

### Immunoprecipitation, immunoblotting and Vangl2 tyrosine phosphorylation

To detect tyrosine phosphorylation of Vangl2, endogenous or exogenous HA- or GFP-tagged Vangl2 proteins were pulled down with rabbit polyclonal Vangl2 (Millipore Sigma, Abn2242), rabbit polyclonal Vangl2 H55 (Santa Cruz), rabbit HA (Bethyl), antibodies and Protein A Sepharose, or GFP- or HA-trap (Chromotek) beads. The analysis of tyrosine phosphorylation was performed using a mouse monoclonal anti-phospho-tyrosine antibody (Santa Cruz, pY20, sc508). For analysis of FGFR1-Vangl2 complexes, FGFR1-3xFLAG RNA was co-injected with GFP-Vangl2 (100 pg each) or PTK7-GFP RNA into four animal blastomeres at four-to-eight cell stage. Embryos were lysed at stage 12 and FGFR1 was pulled down with FLAG beads (Millipore Sigma). Vangl2, PTK7 and FGFR1 were detected in pulldowns and lysates with anti-GFP (GFP-B2, Santa Cruz) or anti FLAG (M2, Millipore Sigma) antibodies. Phosphorylation at specific serine/threonine sites of Vangl2 was analyzed by immunoblotting after pulldowns of HA-Vangl2 with anti-HA antibodies (12CA5), from stage 12 embryo lysates. Membranes were probed with rabbit monoclonal antibodies against phospho-T78/S79/S82 (Cluster 1; AP1206, AbClonal) and phospho-S15/S17 (Cluster 2; AP1204, AbClonal).

The association between endogenous Vangl2 and FGFR1 was analyzed in the neural progenitor cell line C17.2 ^69^ and HEK293T cells. Confluent C17.2 or HEK293T cells, initially plated in 100 mm dishes at 10⁴ cells per cm² were lysed, and immunoprecipitation was performed using 1 µg of either control Protein A Sepharose-purified rabbit IgG or rabbit polyclonal Vangl2 antibody (Millipore Sigma, ABN2242). Pulldowns and lysates were probed with rabbit monoclonal FGFR1 (Cell Signaling, D8E4) and mouse monoclonal Vangl2 (Santa Cruz, C8) antibodies.

### Vangl2 analysis in mouse embryonic stem (ES) cells

Mouse wild type and FGFR1/2 double knockout ES cells ^52^ were cultured in DMEM supplemented with penicillin/streptomycin, 1 mM of sodium pyruvate, non-essential amino acids, 1 µM β-mercaptoethanol, 15% fetal bovine serum (FBS), 1000 U of mouse recombinant LIF (R&D systems), 1 µM of MEK1 inhibitor PD0325901 (Tocris), and 3 µM of GSK3 inhibitor CHIR99021 (Tocris) as described^52^. Before the lysis, cells were plated on gelatinized 60 mm culture dishes. On the next day cells were briefly washed with PBS and cultured for 4 hrs in medium without inhibitors, LIF and FBS, to alleviate potential effects on FGFR signaling. Immunoprecipitation was done with either 1 µg Vangl2 H55 (Santa Cruz) or 0.4 µg of anti-pY antibody (pY20, Santa Cruz) with the lysates from 5×10^6^ cells. Vangl2 pulldowns were immunoblotted with mouse Vangl2-specific antibody (C8, Santa Cruz), pY pulldowns were immunoblotted with rat anti-Vangl2 monoclonal antibody (MABN750, Millipore Sigma) and rabbit Cyclin B1 antibody (H-433, Santa Cruz) that served as negative control. Vangl2 and FGFR1 expression in lysates was analyzed with rat monoclonal Vangl2 (MABN750, Millipore Sigma) and rabbit anti-FGFR1 antibody (D8E4, Cell Signaling) respectively.

## Statistics

Statistical analyses, histograms, and graphs were generated using Prism 10 and Microsoft Excel. A two-tailed Student’s *t*-test (**Fig. 1**, **Fig. S3**, **Fig. S4**) was used to assess statistical significance. Fisher’s exact test was used for comparison of categorical data (**Fig. 6** and **Fig. S1**).

## Data availability

All data used in the analysis are contained within the main text or in the supplementary materials. Data and materials are available on request from the authors.

## ACKNOWLEDGEMENTS

We thank Phil Soriano, Annette Borchers, Miho Matsuda and Jean-Pierre Saint-Jeannet for plasmids, Elena Torban for the anti-Vangl2 antibodies. We are grateful to Phil Soriano for the FGFR1/2 -/- double knockout mouse ES cells, Evan Snyder for C17.2 cells, and Nikos Tzavaras and Shilpa Dilip Kumar from the ISMMS Microscopy Core Facility for cell polarity quantification advice. We also thank James Clark and Jean-Pierre Saint-Jeannet for comments on the manuscript and Sokol lab members for discussions. This research was supported by the NIH grant R35GM122492 to S.Y.S.

## Author contribution statement

I.C. and S.Y.S. initiated, designed the experiments and developed the project. I.C. performed all the experiments, analyzed the data and prepared the figures. I.C. and S.Y.S. wrote the manuscript. S.Y.S. acquired the funding. Both authors approved the final manuscript.

## Supplementary material

**Supplementary Fig. 1.**
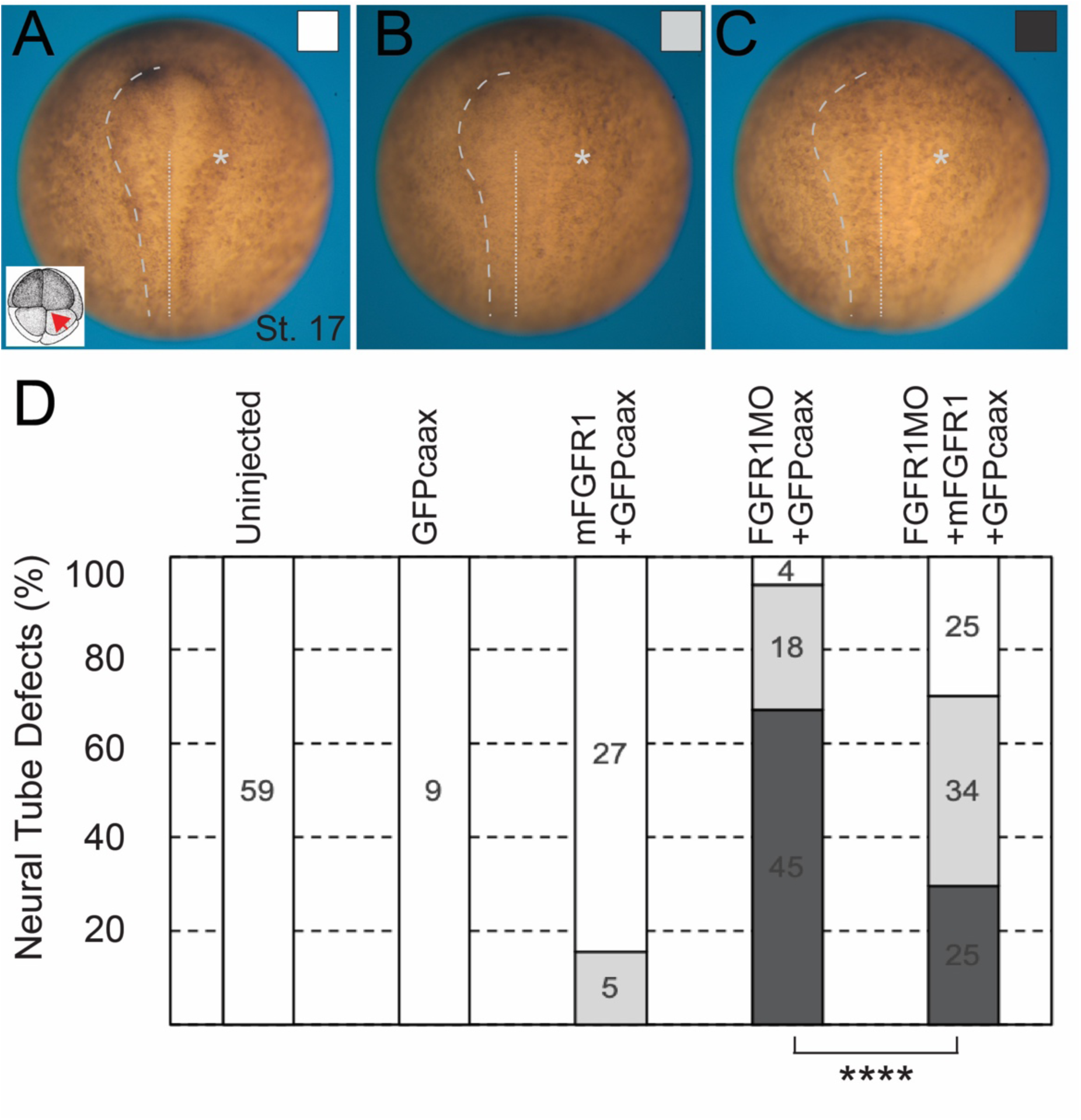
Injection of mouse FGFR1 RNA rescues neural tube defects (NTDs) in FGFR1 morphants. Xenopus embryos were separately injected unilaterally with FGFR1 morpholino (MO, 40 ng) or mouse FGFR1 RNA (60 pg) and GFP-CAAX RNA (200 pg), as lineage tracer, into one dorsal animal blastomere of the 4-8 cell stage. GFP fluorescence was examined at stages 14-15 to confirm correct targeting. Embryos were fixed at stage 16-17, and NTD phenotypes were classified into none (A), mild (B) or severe (C) marked by white, light grey and dark grey squares, respectively, and scored in (D). Position of the neural folds is indicated by dashed lines on the non-injected side. Asterisks mark the injected side. Midline is marked by dotted line. (D) Frequencies of mild and severe NTDs are shown. Numbers of scored embryos per group are indicated. The significance was determined by full 2×3 Fisher’s exact test; **** p<0.0001. Data are representative of three experiments.

**Supplementary Fig. 2.**
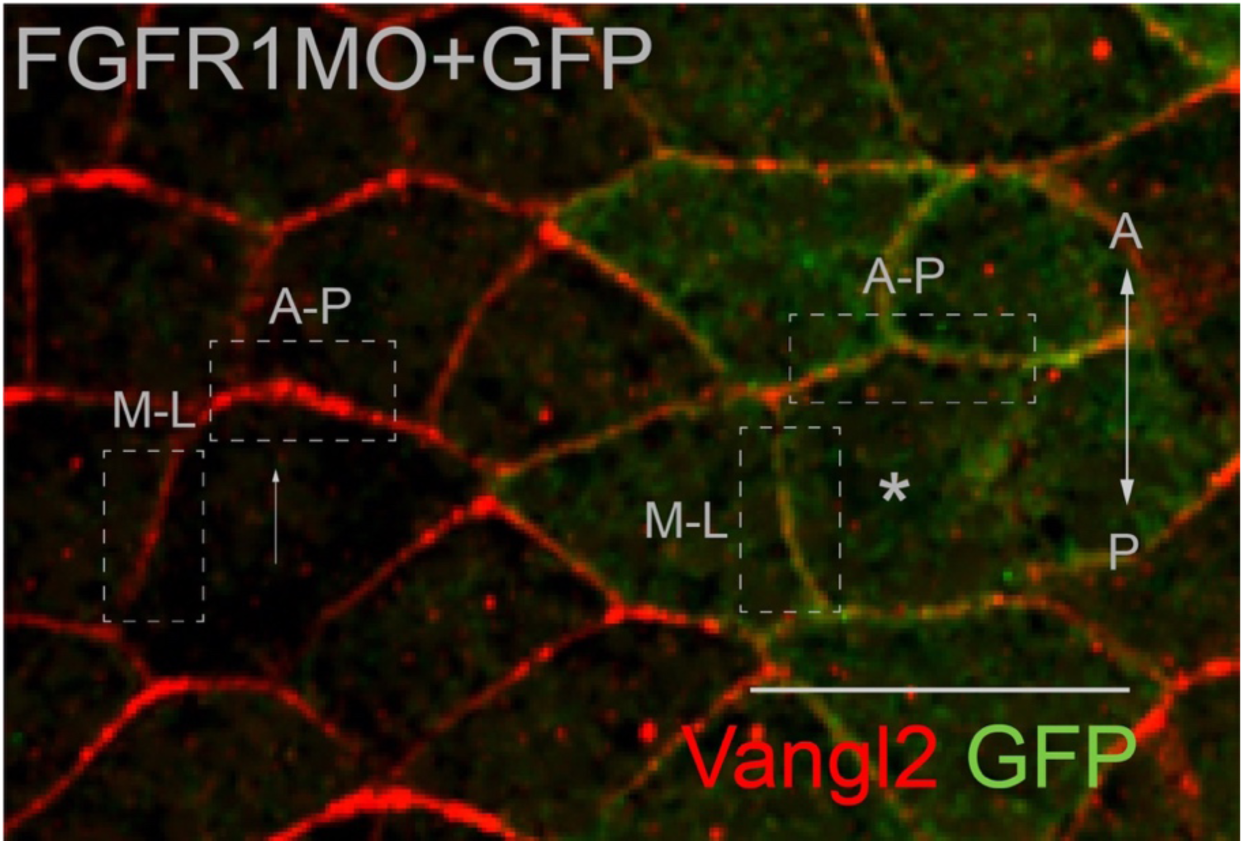
Quantifying planar polarity of neuroepithelial cells (portion of an image shown in Fig. 1C, refers to Fig. 1D). Representative image of neural plate, stage 15 of an embryo injected with FGFR1 morpholino (MO) and GFP RNA, and co-immunostained for Vangl2 and GFP as described in Fig. 1 legend and Methods. A cell was scored as polarized if the fluorescence intensity at an anteroposterior (A-P) junction was at least twofold higher compared to the fluorescence intensity at the mediolateral (M-L) junction. Examples of a planar-polarized cell (arrow) and a non-polarized cell depleted of FGFR1 (asterisk) are shown. Merged channels (Vangl2 (red) +GFP(green)) are shown. A-P and M-L junctions are indicated by dashed boxes. Anteroposterior (A-P) axis is marked. Scale bar: 50 µm.

**Supplementary Fig. 3.**
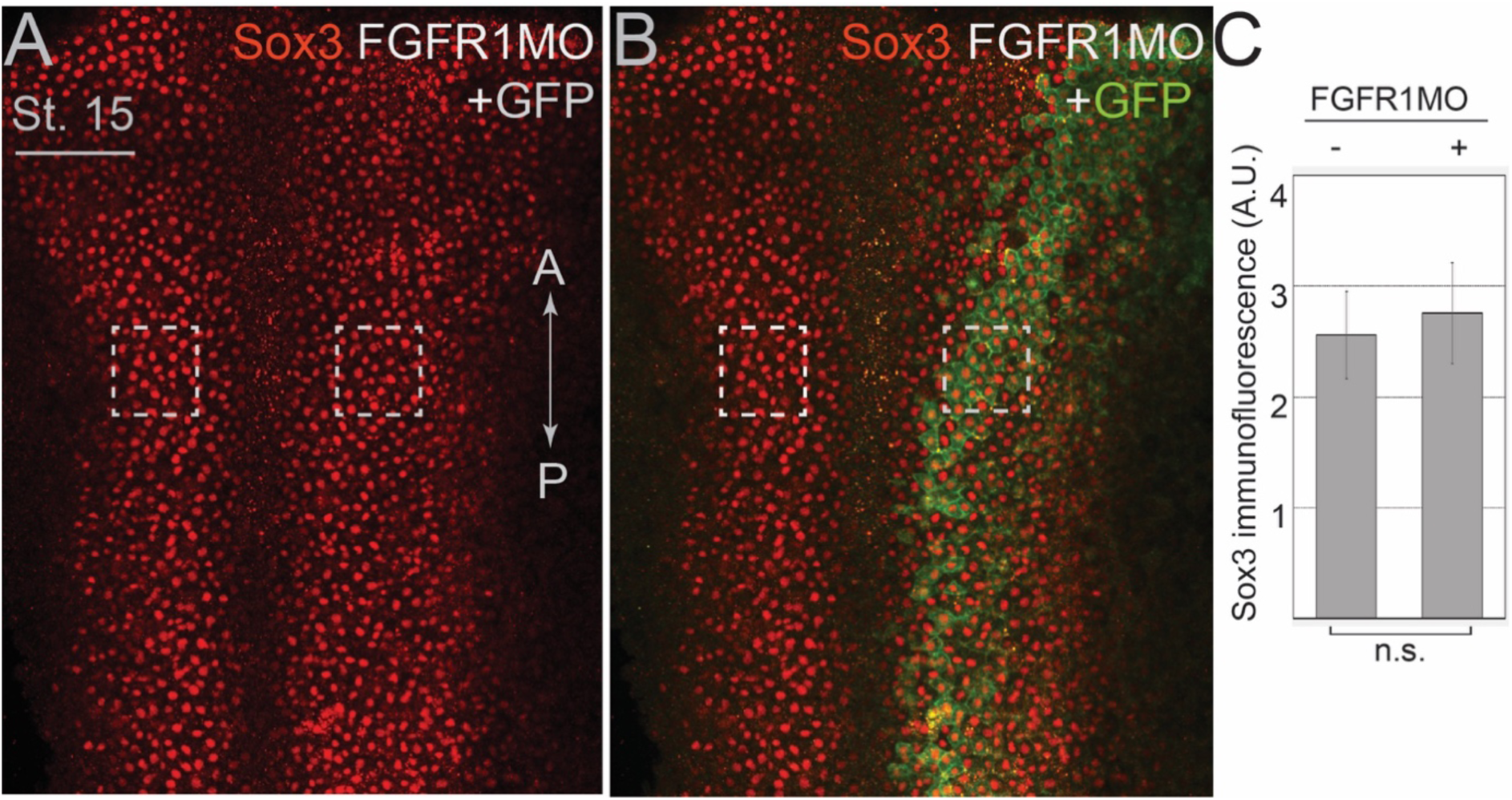
Levels of the pan-neural marker Sox3 are not affected in FGFR1 morphants. Sixteen-cell *Xenopus* embryos were co-injected with 5 nl of FGFR1 MO (15 ng) and GFP RNA (90 pg), as lineage tracer into one dorsal animal blastomere. Embryos were fixed at stage 15 (st.15) and co-immunostained for Sox3 and GFP. Anteroposterior (A-P) axis is indicated. Scale bar: 200 µm. Top view neural plate images show red (Sox3, A) and merged (GFP+Sox3, B) channels. GFP-positive morphant cells retain the expression of Sox3 (dashed squares). (C) Graph showing mean ± s.d. of Sox3 immunofluorescence (A.U., arbitrary units) in uninjected and FGFR1MO+GFP-injected neuroepithelium. Fluorescence was quantified in 3 × 10⁴ µm² areas containing 60–90 cells per square in three independent embryos. Statistical significance was assessed using a two-tailed Student’s *t*-test (n.s., not significant). Data represent two independent experiments.

**Supplementary Fig. 4.**
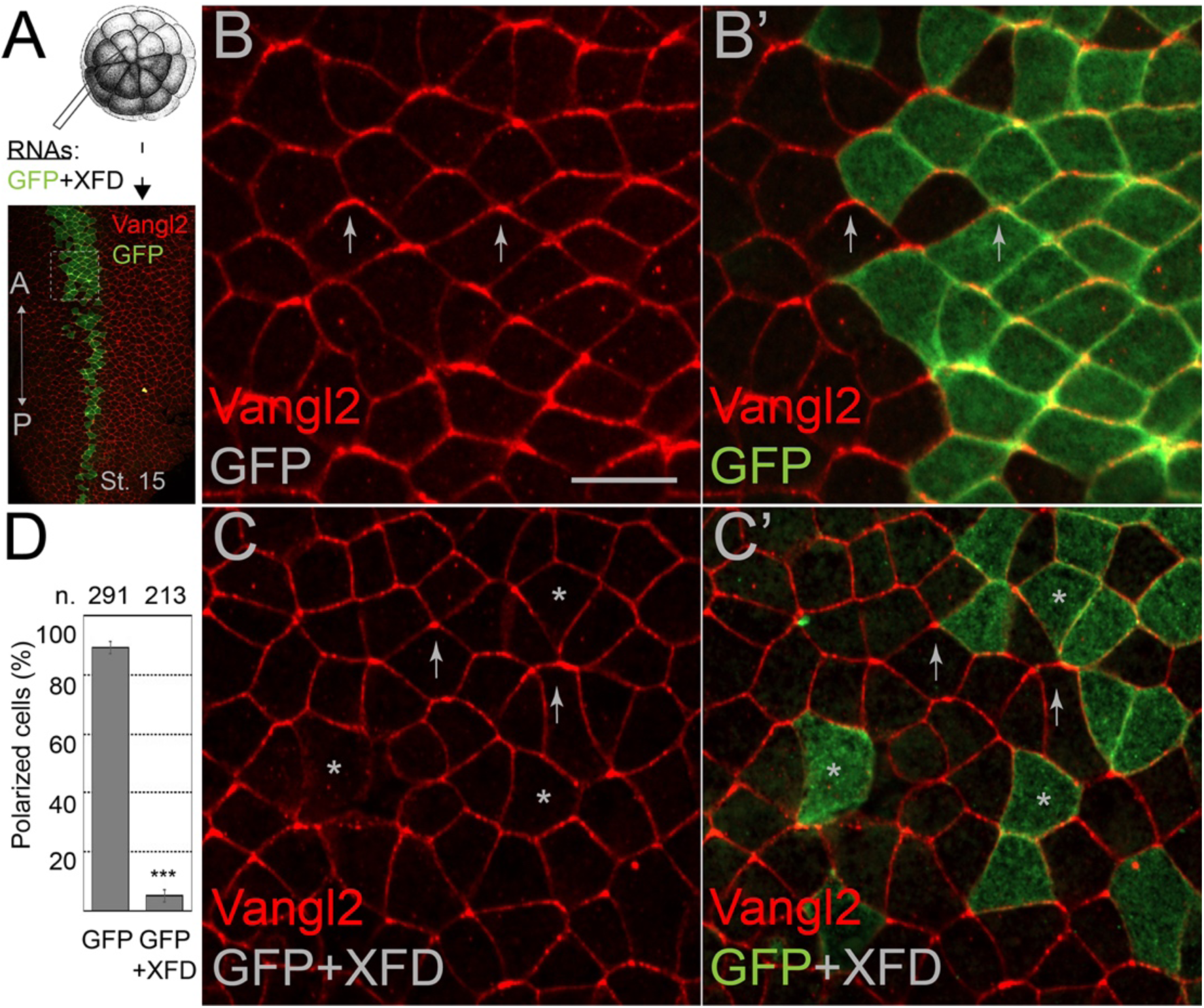
Dominant-interfering FGFR1 construct blocks the neural plate PCP. (A) Experimental scheme. Thirty-two-cell stage *Xenopus* embryos were injected with RNAs encoding GFP (200 pg) and dominant-interfering FGFR1 construct (XFD, 400 pg) into one dorsal animal blastomere. Embryos were collected at stage 15 and co-immunostained for Vangl2 (red) and GFP (green). The dashed boxed area is magnified in (B-C’). The anteroposterior (A-P) axis is indicated. (B-B’) Representative images of neural plate cells expressing GFP tracer with anteriorly enriched Vangl2 (arrows). (C-C’) Vangl2 is not restricted to anterior cell boundaries in cells expressing XFD (asterisks). (D) Quantification of the mean ± s.d. frequencies of cells with anteriorly enriched Vangl2. Numbers of scored cells (n) are indicated on top of each bar. Three embryos were scored for each group (40-100 cells per embryo). Data represent three independent experiments. Statistical significance was assessed using a two-tailed Student’s *t*-test (***, p<0.001). Scale bar: 30 µm.

**Supplementary Fig. 5.**
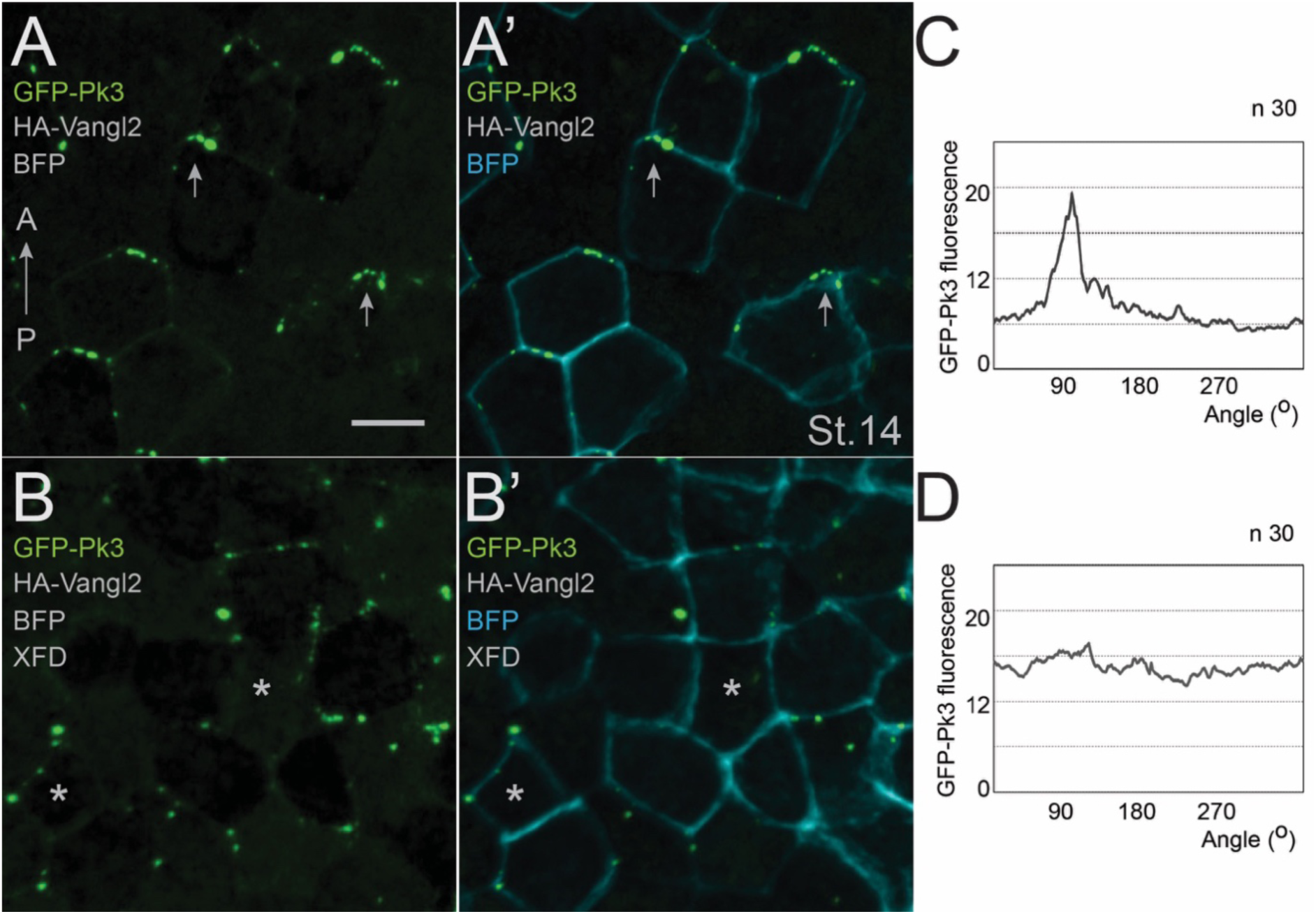
Dominant-interfering FGFR1 construct inhibits the anterior recruitment of Pk3 complexes in the neural plate. Thirty-two-cell stage *Xenopus* embryos were injected into one dorsal animal blastomere with RNAs encoding GFP-Pk3 (150 pg), HA-Vangl2 (20 pg), and myristoylated BFP (BFP, 80 pg), with or without the dominant-interfering FGFR1 construct (XFD, 400 pg), and fixed at stage 14 (st.14). GFP-Pk3 (Pk3, green) and BFP (blue) fluorescence channels are shown. GFP-Pk3 (A, B) and merged GFP-Pk3+BFP fluorescence (A’, B’) is shown. Anteroposterior (A-P) axis is indicated. Dorsal view of representative images of neural plate cells expressing GFP-Pk3 without (A-A’, arrows) and with XFD (B-B’, asterisks). Cell membranes are marked by BFP (A’, B’). Scale bar: 30 µm. (C, D) GFP-Pk3 fluorescence around cell boundaries in control (C) and XFD-expressing (D) mosaic neuroepithelial cells was quantified as described in Figure 5 legend and Methods and plotted as a function of angle. In control cells, GFP-Pk3 fluorescence is enriched at the anterior side (90°), whereas it is evenly distributed in cells co-expressing XFD. Cell numbers (n) are shown.

**Supplementary Fig. 6.**
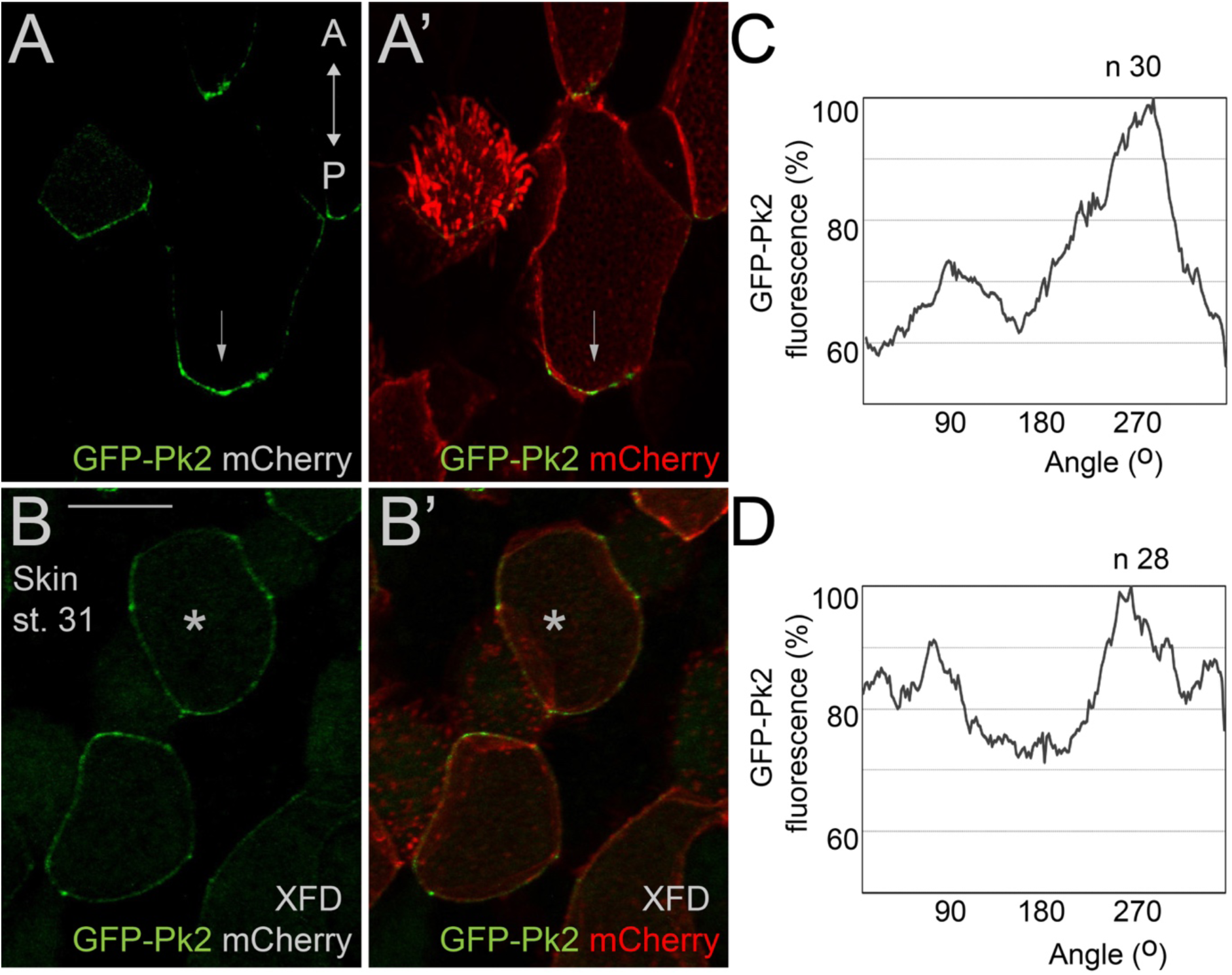
Dominant interfering FGFR construct (XFD) inhibits PCP in *Xenopus* skin. Thirty-two-cell stage embryos were injected into one ventral blastomere with RNAs encoding GFP-Pk2 (700 pg) and mCherry (100 pg), with or without XFD (2 ng). Late tailbuds, stage 31 (st. 31) were fixed and the polarity of the PCP marker GFP-Pk2 was assessed in mosaic skin cells. GFP-Pk2 is polarized in control cells (A-A’, arrows), but not in embryos co-injected with XFD (B-B’, asterisks). Single-channel images for GFP (A, B) and images showing merged GFP+mCherry channels (A’, B’), are shown. Double-headed arrow indicates the anteroposterior (A-P) axis. Scale bar: 30 µm. (C, D) Average GFP-Pk2 fluorescence around the cell perimeter was quantified in Fiji and plotted as a function of angle (degrees, °), expressed as percentiles (%). The posterior side corresponds to 270°. GFP-Pk2 is posteriorly enriched in control embryos (C), but not in those co-injected with XFD (D). A total of 30 and 28 cells (n) were quantified in (C) and (D), respectively.

**Supplementary Fig. 7.**
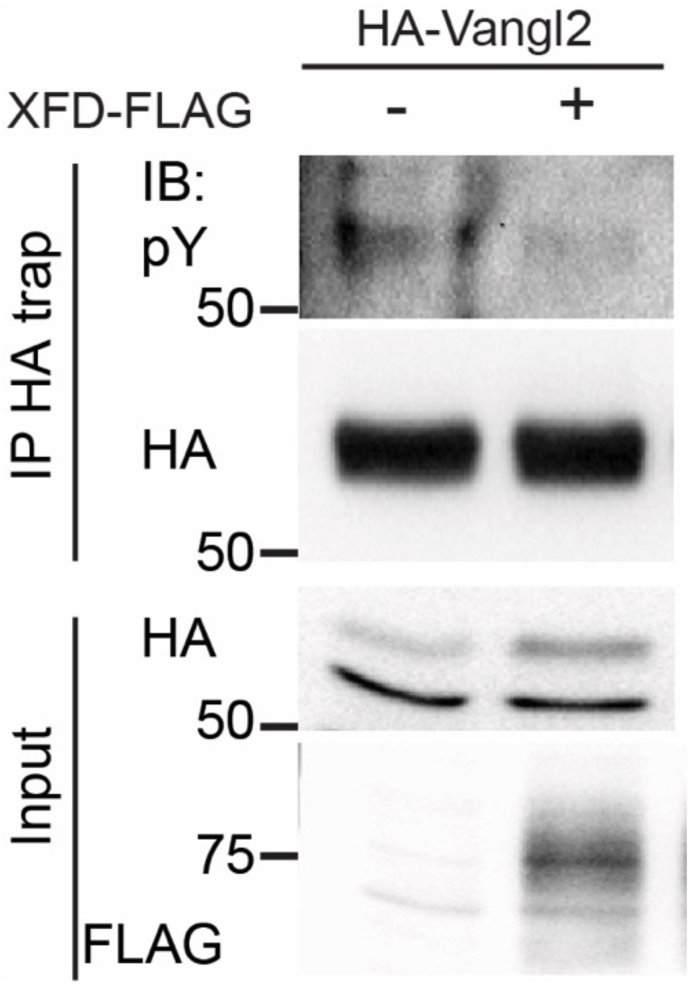
Dominant-interfering FGFR1-FLAG inhibits Vangl2 tyrosine phosphorylation. *Xenopus* embryos were injected with RNAs for HA-Vangl2 (40 pg) with or without XFD-3xFLAG (150 pg) and cultured until stage 12. Vangl2 tyrosine phosphorylation was analyzed by HA-trap pulldown followed by immunoblotting (IB) with anti-pY, anti-HA, or anti-FLAG antibodies as indicated.

**Supplementary Fig. 8.**
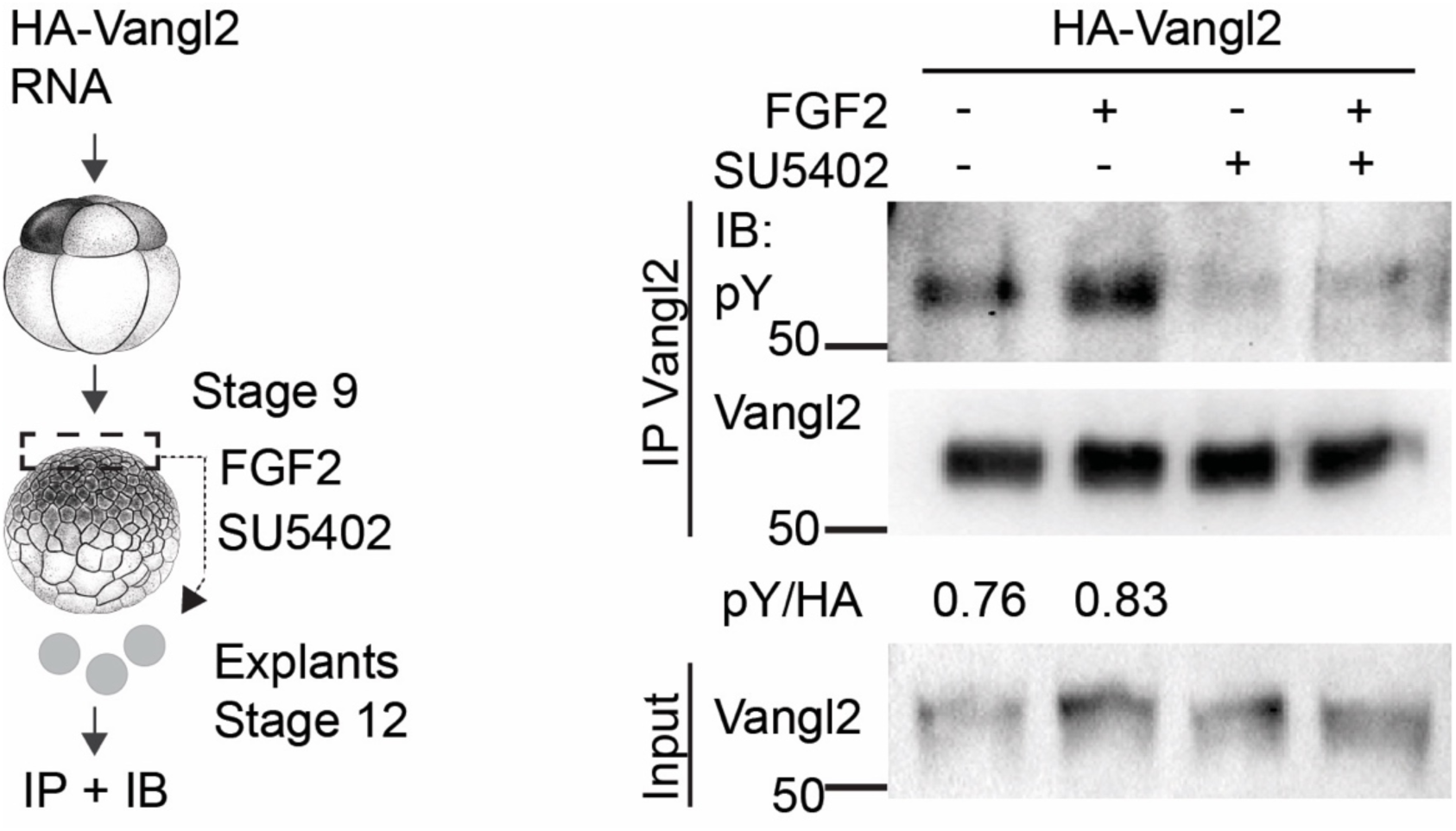
Inhibition of endogenous FGFR signaling with SU5402 reduces Vangl2 tyrosine phosphorylation. Left: Experimental scheme. Ectoderm explants were dissected from *Xenopus* embryos injected with HA-Vangl2 RNA (40 pg) at stage 9 and cultured in the presence of FGF2 (25–100 ng/ml), with or without SU5402 (100 µM), until stage 12. Vangl2 phosphorylation was analyzed in anti-Vangl2 immunoprecipitates immunoblotted with anti-phosphotyrosine (pY) and anti-HA antibodies. Intensity ratios of pY to HA signals for control and FGF2-treated explants are shown.

**Supplementary Fig. 9.**
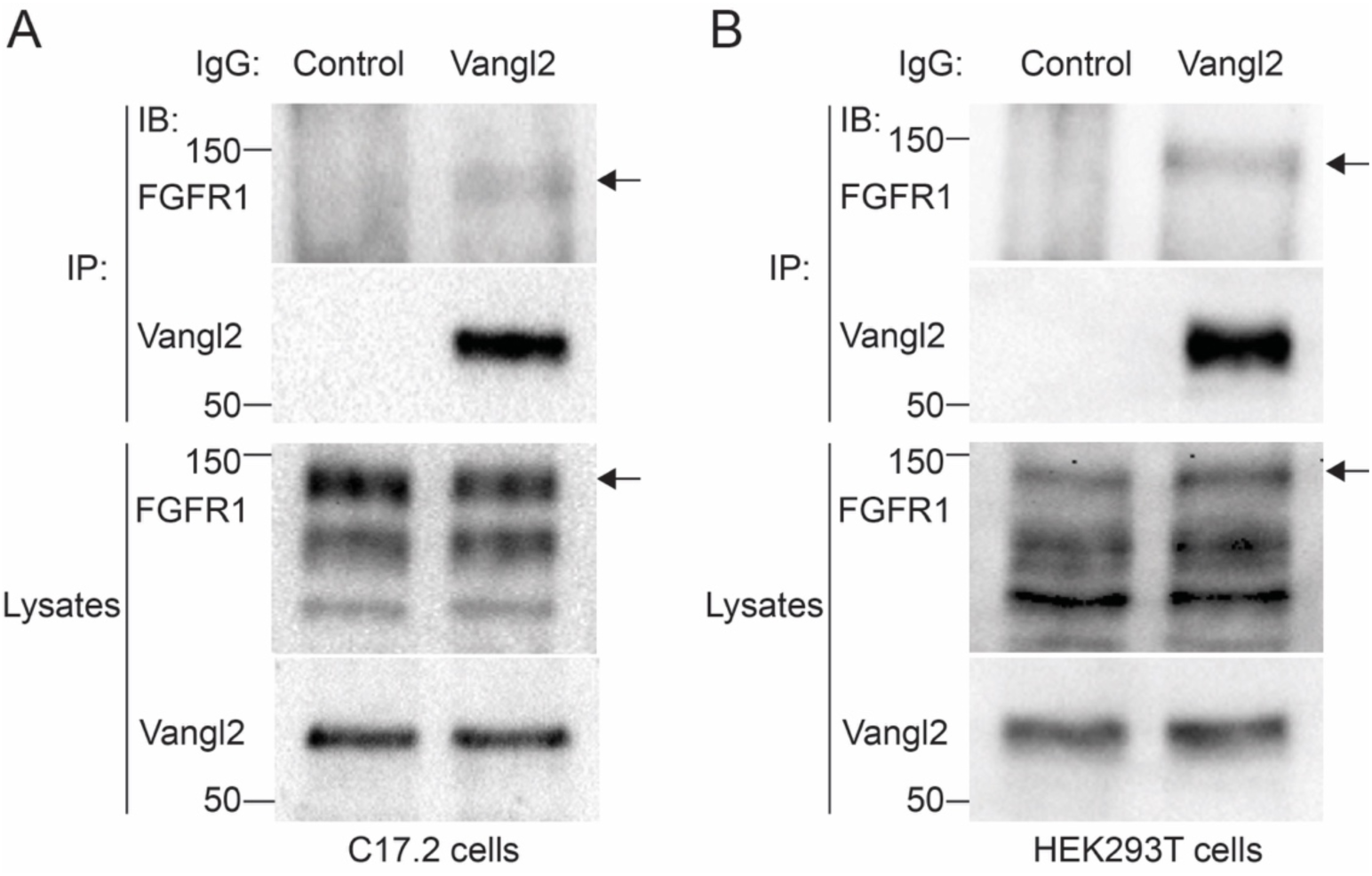
Physical association between endogenous Vangl2 and FGFR1. Formation of the complex between Vangl2 and FGFR1 was assessed in C17.2 neural progenitor cells (A) and HEK293T (B) cells. Cell lysates were precipitated with 1 µg of anti-Vangl2 antibody or control rabbit IgG. The immunoprecipitates and input lysates were probed with FGFR1 and Vangl2 antibodies. The band corresponding to the upper FGFR1 band in C17.2 and HEK293T cell lysates is detected in Vangl2 pulldowns from both cell lines (A, B; arrows). Molecular weight markers are indicated.

**Supplementary Fig. 10.**
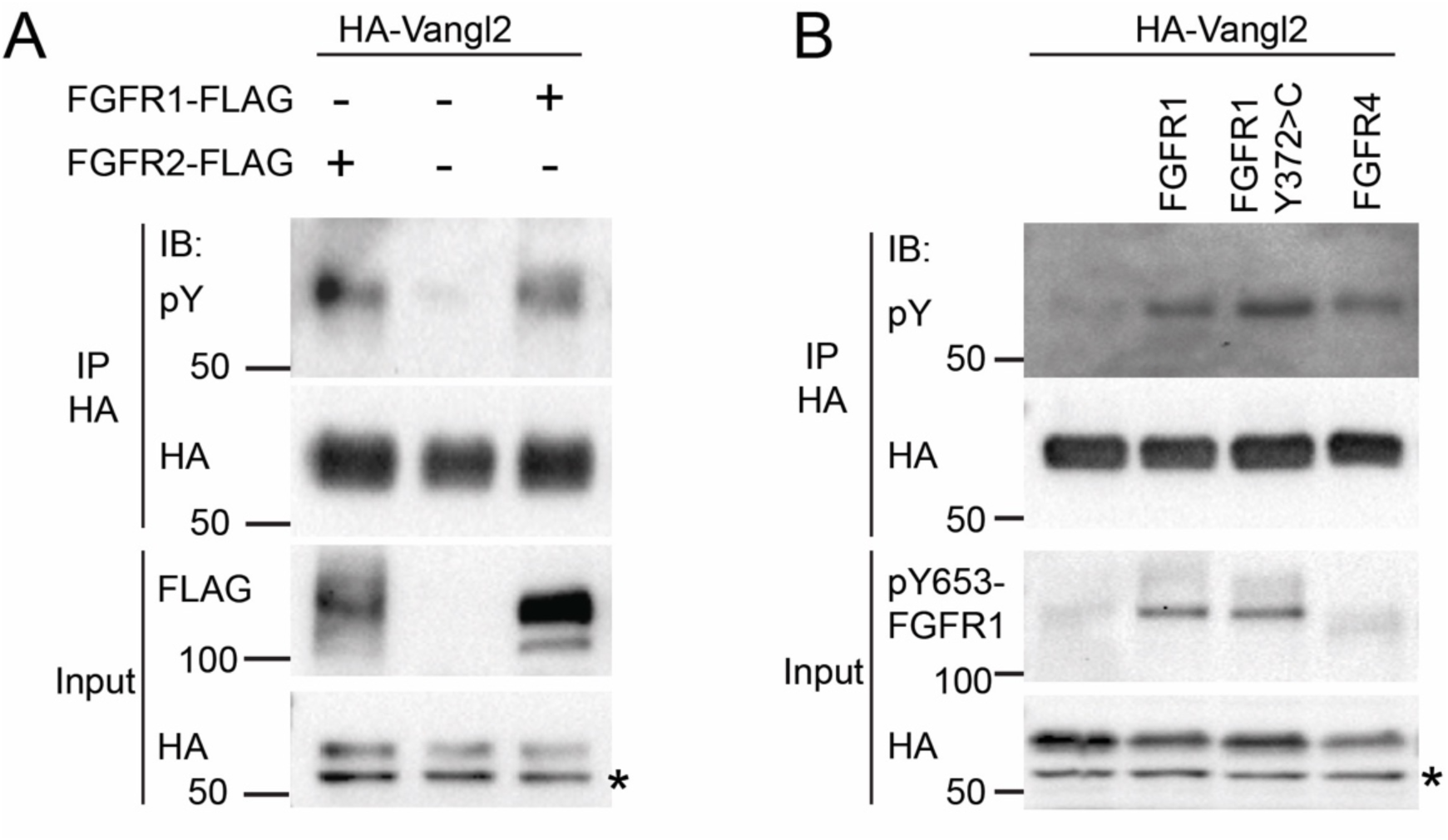
Induction of Vangl2 tyrosine phosphorylation by FGF receptors. (A) *Xenopus* embryos were injected with RNAs for Xenopus HA-Vangl2 (40 pg) with or without mouse FGFR1 and FGFR2 (40 pg each), or (B) DNA constructs encoding HA-Vangl2 (25 pg) with or without human FGFR1, FGFR1 Y372>C, or FGFR4 (50 pg each), (see Methods), and cultured until stage 12 (A) or stage 14 (B). Vangl2 tyrosine phosphorylation (Vangl2-pY) was analyzed by pull-downs with rabbit anti-HA antibody (A) or HA-trap beads (B), immunoblotted with anti-pY, anti-HA, or anti-FLAG antibodies as indicated. Expression of wild-type and mutated human FGFR constructs (B) was assessed with anti-phospho-Y653-FGFR1 antibody. Asterisk marks non-specific band in lysates (A, B).

**Supplementary Fig. 11.**
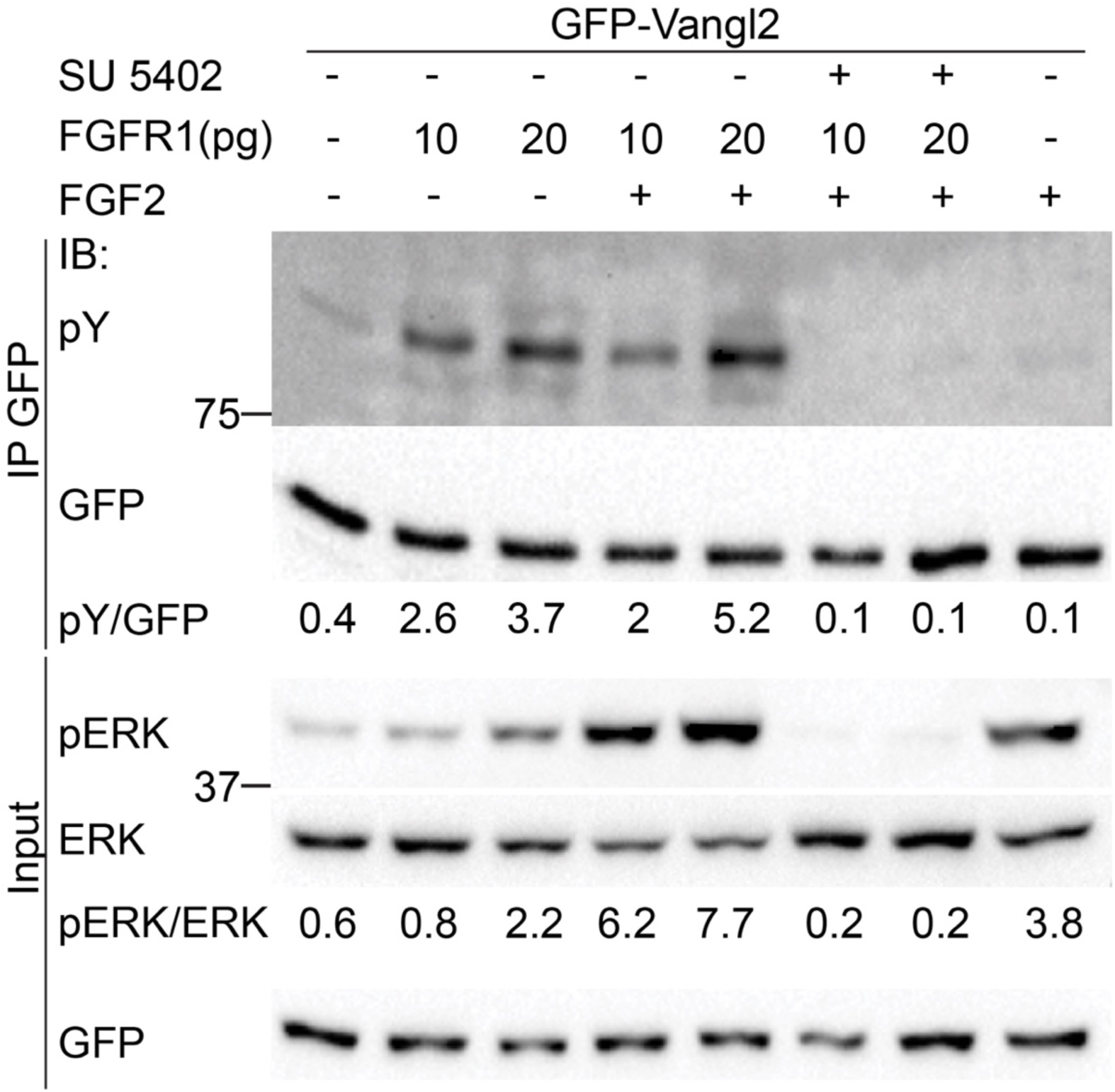
Lack of correlation between Vangl2 phosphorylation and ERK activation in response to FGFR1 signaling. Four- to eight-cell *Xenopus* embryos were injected into four animal blastomeres with GFP-Vangl2 RNA (40 pg), with or without FGFR1-FLAG RNA (10 pg or 20 pg). Ectoderm explants were dissected at stage 9, cultured until stage 12, lysed, and GFP-Vangl2 was pulled down using GFP-trap beads. Ectoderm explants were cultured with or without FGF2 (100 ng/mL) or SU5402 (100 µM). FGFR1 dose-dependently induced Vangl2 tyrosine phosphorylation. FGF2 strongly activated ERK (pERK) and weakly induced Vangl2 phosphorylation. Neither Vangl2 nor ERK phosphorylation was detected in SU5402-treated explants. Data represent three independent experiments.

**Supplementary Fig. 12.**
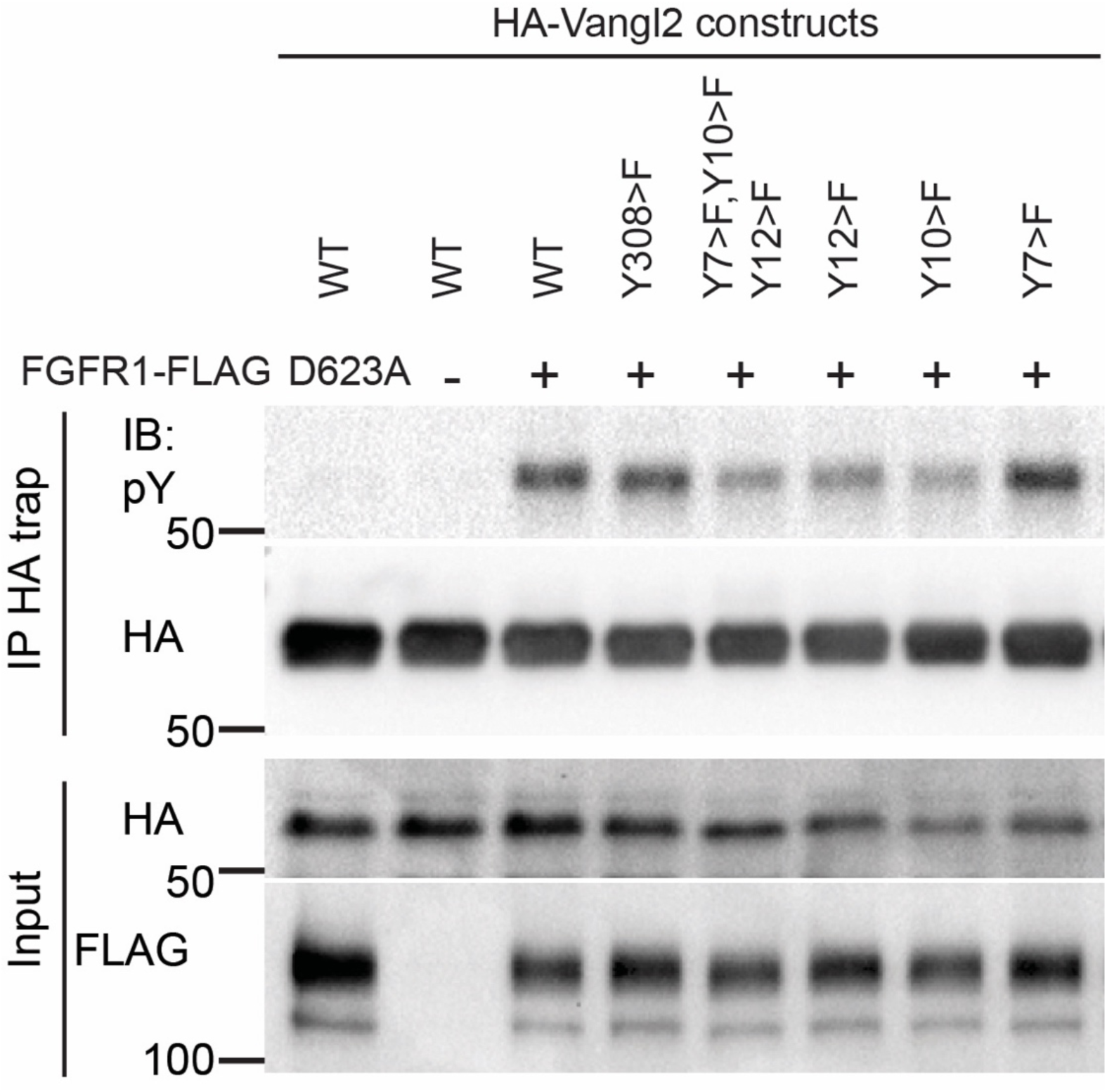
Mapping major FGFR1-dependent phospho-tyrosine sites in Vangl2. Embryos were injected with 40 pg of RNAs encoding FGFR1, FGFR1D623A and various HA-Vangl2 constructs as indicated. Vangl2 phosphorylation was analyzed in HA-trap immunoprecipitates from stage 12 embryo lysates. Immunoblotting (IB) was done with anti-pY, anti-HA or anti-FLAG antibodies as indicated.

**Supplementary Fig. 13.**
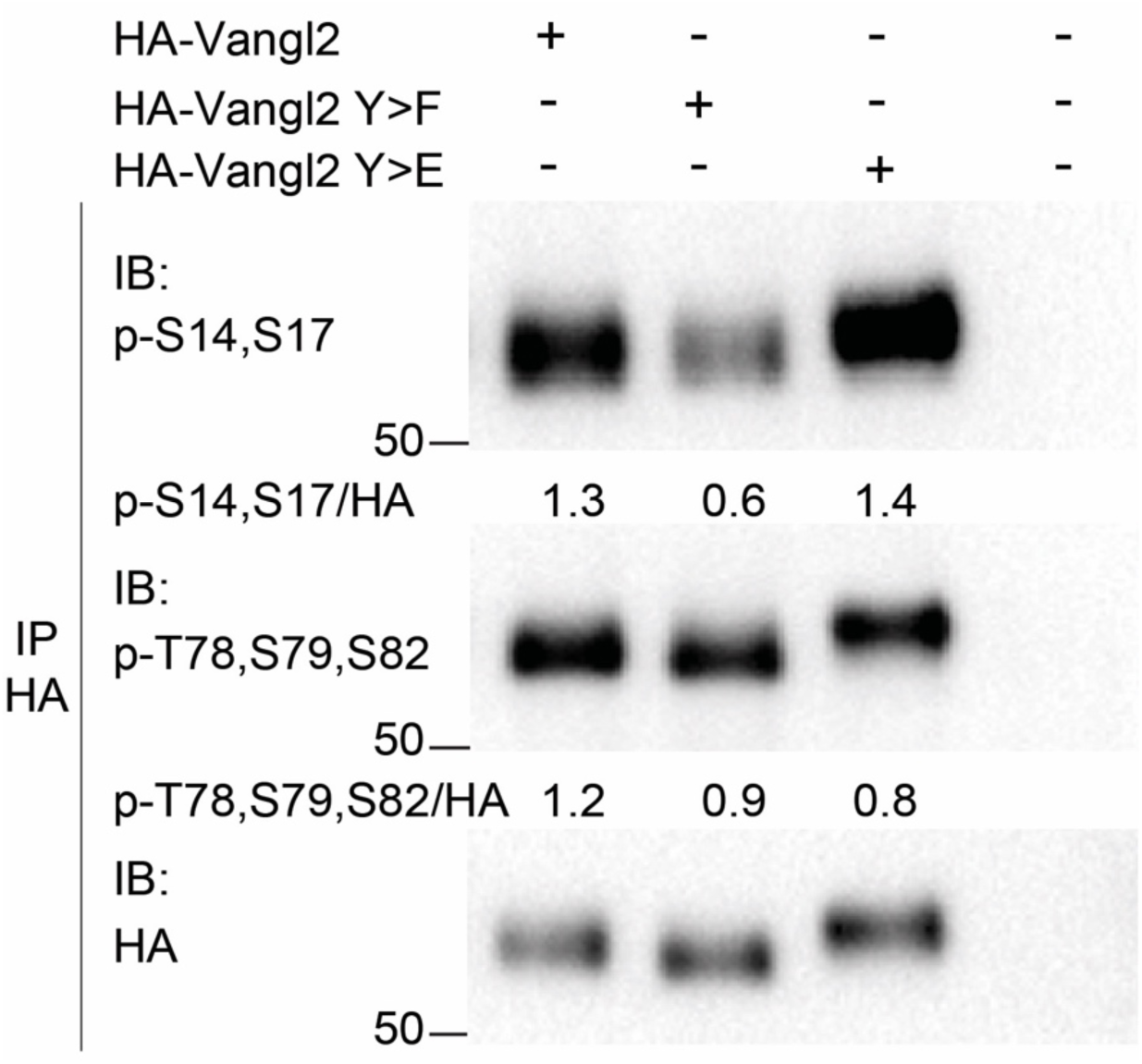
Inhibition of Cluster II serine phosphorylation in Vangl2 construct with the substituted tyrosine phosphorylation sites. Four- to eight-cell *Xenopus* embryos were injected with RNAs encoding HA-Vangl2, Y7Y10Y12>F (Y>F), or Y7Y10Y12>E (Y>E) constructs (40 pg each). Vangl2 proteins were pulled down from stage 13 lysates using anti-HA antibody and probed with anti-pS14S17, anti-pT78S79S82, and anti-HA antibodies. Band intensity ratios (pS14S17/HA and pT78S79S82/HA) are shown. Molecular weights are indicated. Data represent two independent experiments.

